# Domain topology, stability, and translation speed determine mechanical force generation on the ribosome

**DOI:** 10.1101/490821

**Authors:** Sarah E. Leininge, Fabio Trovato, Daniel A. Nissley, Edward P. O’Brien

## Abstract

The concomitant folding of a nascent protein domain with its synthesis can generate mechanical forces that act on the ribosome and alter translation speed. Such changes in speed can affect the structure and function of the newly synthesized protein as well as cellular phenotype. The domain properties that govern force generation have yet to be identified and understood, and the influence of translation speed is unknown as all reported measurements have been carried out on arrested ribosomes. Here, using coarse-grained molecular simulations and statistical mechanical modeling of protein synthesis, we demonstrate that force generation is determined by a domain’s stability and topology, as well as translation speed. The statistical mechanical models we create predict how force profiles depend on these properties. These results indicate that force measurements on arrested ribosomes will not always accurately reflect what happens in a cell, especially for slow-folding domains, and suggest the possibility that certain domain properties may be enriched or depleted across the structural proteome of organisms through evolutionary selection pressures to modulate protein synthesis speed and post-translational protein behavior.

**Significance Statement:** Mechanochemistry, the influence of molecular-scale mechanical forces on chemical processes, can occur on actively translating ribosomes through the force-generating actions of motor proteins and the co-translational folding of domains. Such forces are transmitted to the ribosome’s catalytic core and alter rates of protein synthesis; representing a form of mechanical allosteric communication. These changes in translation-elongation kinetics are biologically important because they can influence protein structure, function, and localization within a cell. Many fundamental questions are unresolved concerning the properties of protein domains that determine mechanical force generation, the effect of translation speed on this force, and exactly how, at the molecular level, force is generated. In this study we answer these questions using cutting-edge molecular simulations and statistical mechanical modeling.

---

Mechanical forces acting on a nascent polypeptide chain can alter the speed of protein synthesis by the ribosome^1–7^. Such changes in translation speed can influence the timing and efficiency of a variety of co-translational processes affecting the fold^8,9^, function^10,11^, and intracellular localization of the newly synthesized protein^12,13^. One source of mechanical force is the co-translational folding of individual nascent chain domains as they emerge from the ribosome exit tunnel^1^. The process of folding is presumed to generate a pulling force on residues remaining in the exit tunnel (Fig. 1a,b). That pulling force is transmitted 10 nm back to the catalytic core of the ribosome in a form of mechanical allosteric communication, altering the energy barrier to peptide bond formation^7^, thereby influencing translation speed. The factors governing co-translational force generation have not yet been identified, but they are important to understand as they can modulate translation speed which can alter protein structure and affect the evolutionary selection pressures shaping the sequence and properties of both mRNA and proteins.

**Figure 1.**
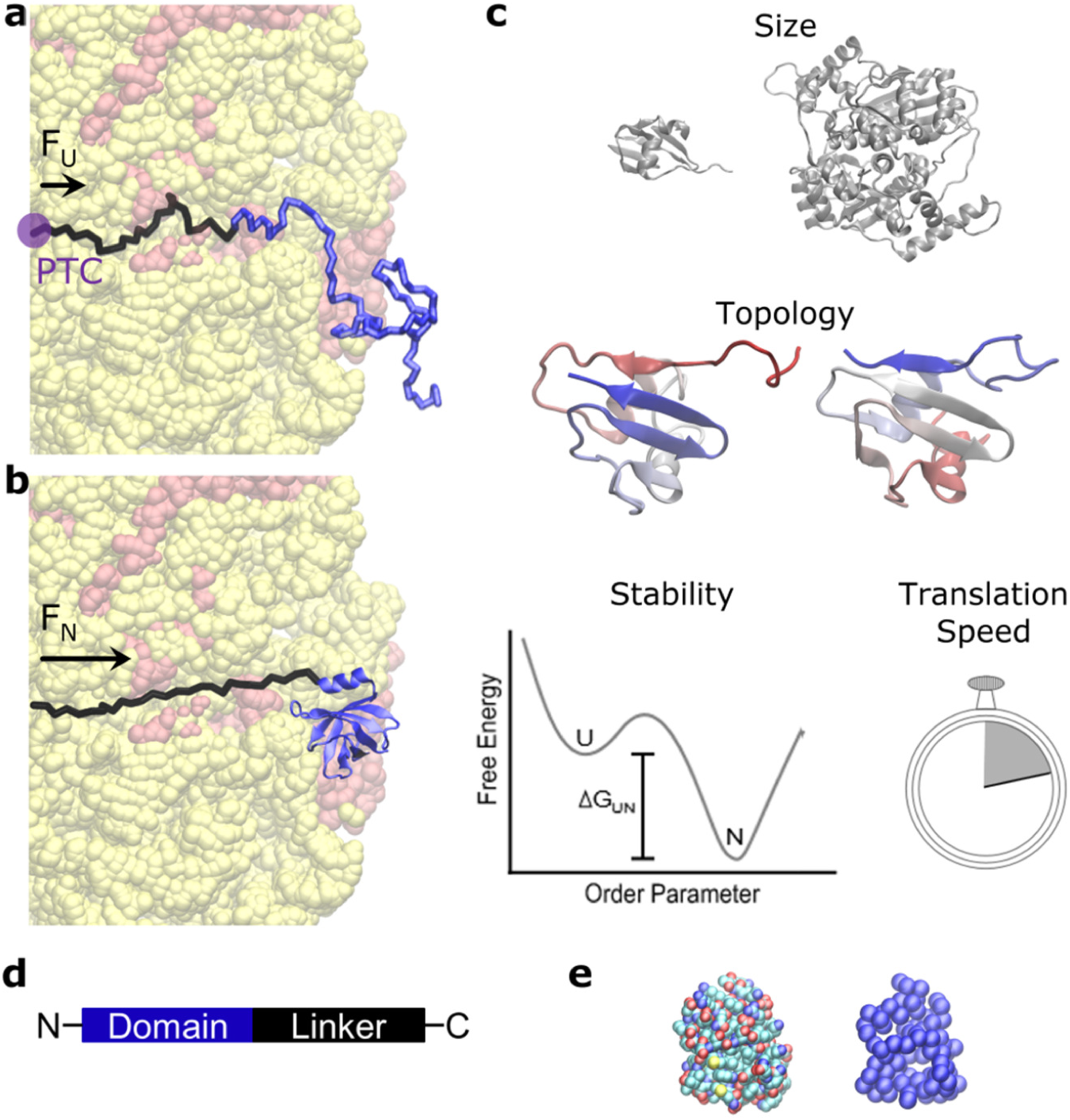
Cotranslational folding and the factors that might govern mechanical force generation. Cotranslational folding of a nascent protein domain (blue, in panels **a** and **b**) on the ribosome (red and yellow) has been implicated in generating a pulling force that is transmitted 10 nm through the exit tunnel to the peptidyl-transferase center. The domain’s unfolded **(a)** and folded state **(b)** each generate a pulling force that is transmitted 10 nm to the PTC of the ribosome and are denoted F_U_ and F_N_, respectively. The difference between these two forces is the force generated by the act of domain folding (Eq. 6). Four factors may influence the extent of force generation: domain size, topology, stability, and translation speed (**c**). Differences in domain size are illustrated by a large and small domain, in gray. Differences in topology are illustrated by the rewiring of unstructured loops between secondary structural elements. (*N.B.*, the colors of the domains shown under ‘Topology’ are from N-terminus to C-terminus, red to blue.) Native state stability is illustrated as a free energy difference, ΔG_UN_, between the unfolded (U) and folded (N) states in the free-energy profile as a function of a hypothetical order parameter. And translation speed is illustrated as a stop watch, to metaphorically time how quickly ribosomes synthesize a protein. **(d)** Each domain in this study has an unstructured linker attached to its C-terminus. **(e)** Illustration of an all-atom (left) and coarse-grained representation (right) of a protein domain.

Goldman and co-workers estimate 12 pN of force is generated when the 106-residue long Top7 protein folds on a translationally arrested ribosome^1^. This force was estimated through a combination of experiments in which Top7’s midpoint folding force was measured off the ribosome, and the force required to relieve stalling by the arrest peptide SecM was measured in the absence of a folding domain^1^. The relative strength of pulling forces generated by different proteins has also been measured using an expression assay comparing the amount of full-length protein produced to the amount of truncated protein produced due to translation stalling by SecM^3^. If the pulling force is large enough at a given nascent chain length it will overcome SecM-induced stalling and produce a full-length protein. Recent papers describe how forces measured in coarse-grained simulations can be used to generate estimates of the fraction of full-length protein^14,15^. Taken together, these studies demonstrate that different proteins produce different forces upon folding that can in turn alter translation speed.

Here, we seek to identify what domain properties determine the magnitude of this pulling force, and to test whether the forces measured in the artificial situation of translationally arrested ribosomes are the same as those that arise during continuous synthesis (Fig. 1c). To do this, we simulate the synthesis of five different *E. coli* proteins by the *E. coli* ribosome using coarse-grained models. We find that three factors, the domain’s stability, topology, and translation rate, are the primary determinants of force generation, and identify the conditions under which the forces measured on arrested ribosomes do not correspond to those measured on continually translating ribosomes. We also create a non-equilibrium statistical mechanical model that describes how the pulling force changes with codon translation speed, domain stability, and nascent chain length.

## Results

### Force is generated by folding

To examine whether folding generates a force on the catalytic core of the ribosome and to determine its range we ran translationally arrested ribosome-nascent chain simulations at different nascent chain lengths for five different *E. coli* proteins – Protein thiS (PDB ID: 1F0Z)^16^, Pseudouridine synthase D (2IST)^17^, Signal transduction protein PmrD (2JSO)^18^, Protein yrbA (1NY8)^19^ and Protein ybcJ (1P9K)^20^. These proteins are in the α/β structural class and the simulated domains were assigned native-state stabilities typical of those found in experiment (Fig. 2)^21^. An unstructured linker was attached to the C-terminii of the domains to mimic domain folding in the context of a multidomain protein (Fig. 1d). We define linker residues as those nascent chain residues located after the C-terminal residue of the domain because they covalently ‘link’ the folding domain to the P-site tRNA on the ribosome. The linker length is the number of residues composing the linker. The linker lengths used in these simulations were chosen such that they bracketed the length at which the midpoint of folding occurred. It was recently found that the unfolded state of a nascent chain can generate an entropic pulling force on the ribosome^7^, and attractive interactions between the nascent chain and outer ribosome surface also have the potential to give rise to forces^22^. To isolate the force arising from domain folding we ran additional simulations in which the domains always remain unfolded (see SI Appendix) and simulated these folding-incompetent proteins on arrested ribosomes at the same linker lengths. The average pulling force at each linker length was then calculated as the average force at the C-terminus of the nascent chain, which is located at the P-site of the ribosome, minus the average force arising from the folding-incompetent protein (Eq. 6 in Methods).

**Figure 2.**
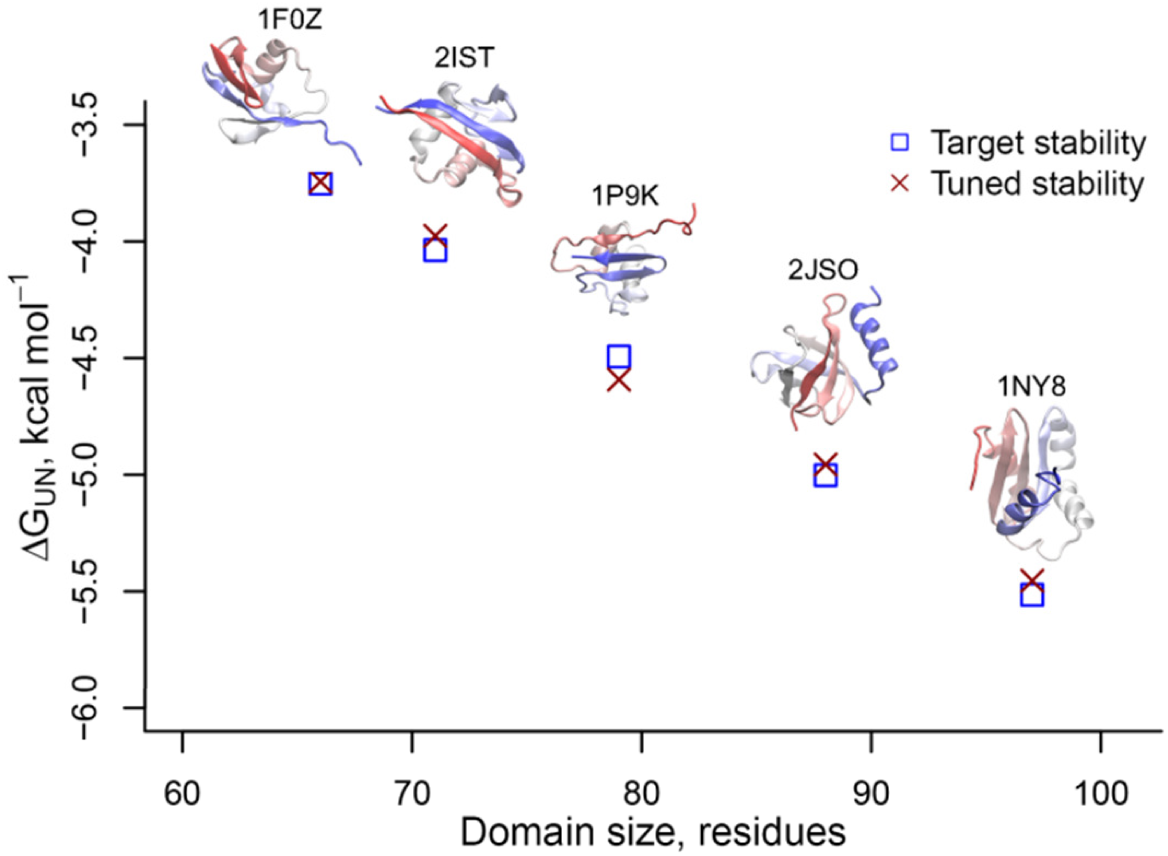
Setting a realistic energy scale for the domain’s coarse-grained force field. The freeenergy difference between the native and unfolded state, Δ*G*_UN_ = -*RT* ln[*P*(*N*)/*P*(*U*)], as a function of domain size, in units of residues, at 310 K, where *R* is the gas constant, *T*, the temperature, and *P*(*N*) and *P*(*U*), the probabilities of being folded and unfolded, respectively. The target stabilities from the PREFUR algorithm^21^ are shown as blue squares, and the free energy of each domain in the tuned simulation force-field are shown as red X’s. The error bars (95% confidence intervals about the mean calculated from Block Averaging) for the calculated free energies are smaller than the size of the symbols. Stabilities are for five different proteins—1F0Z, 2IST, 1P9K, 2JSO, 1NY8, as labeled. Their corresponding crystal structures are displayed in secondary structure format.

For all five proteins we observe that the average pulling force due to folding starts out at zero at short linker lengths, reaches a maximum near the midpoint of the folding transition of the domain, and decreases to zero again after the nascent chain elongates by just a few more residues (Fig. 3). For example, the pulling force for protein 1F0Z is at or near 0 pN below a linker length of 20 residues (blue circles, Fig. 3a), increases to a maximum of 2.5 pN at 22 residues, and then decreases back to 0 pN as the linker length increases to 29 residues. The force maximum at a linker length of 22 residues corresponds to the midpoint of folding as indicated by the inflection point in the average root-mean-square deviation (RMSD) of the simulation structures from the crystal structure versus linker length profile (red squares in Fig. 3a). For these five proteins, we find the maximum folding force ranges between 1.0 and 8.4 pN at linker lengths between 16 and 26 residues and that the linker length locations of maximum force occur at or just after the greatest decrease in the RMSD from the crystal structure, *i.e.*, the midpoint of folding.

**Figure 3.**
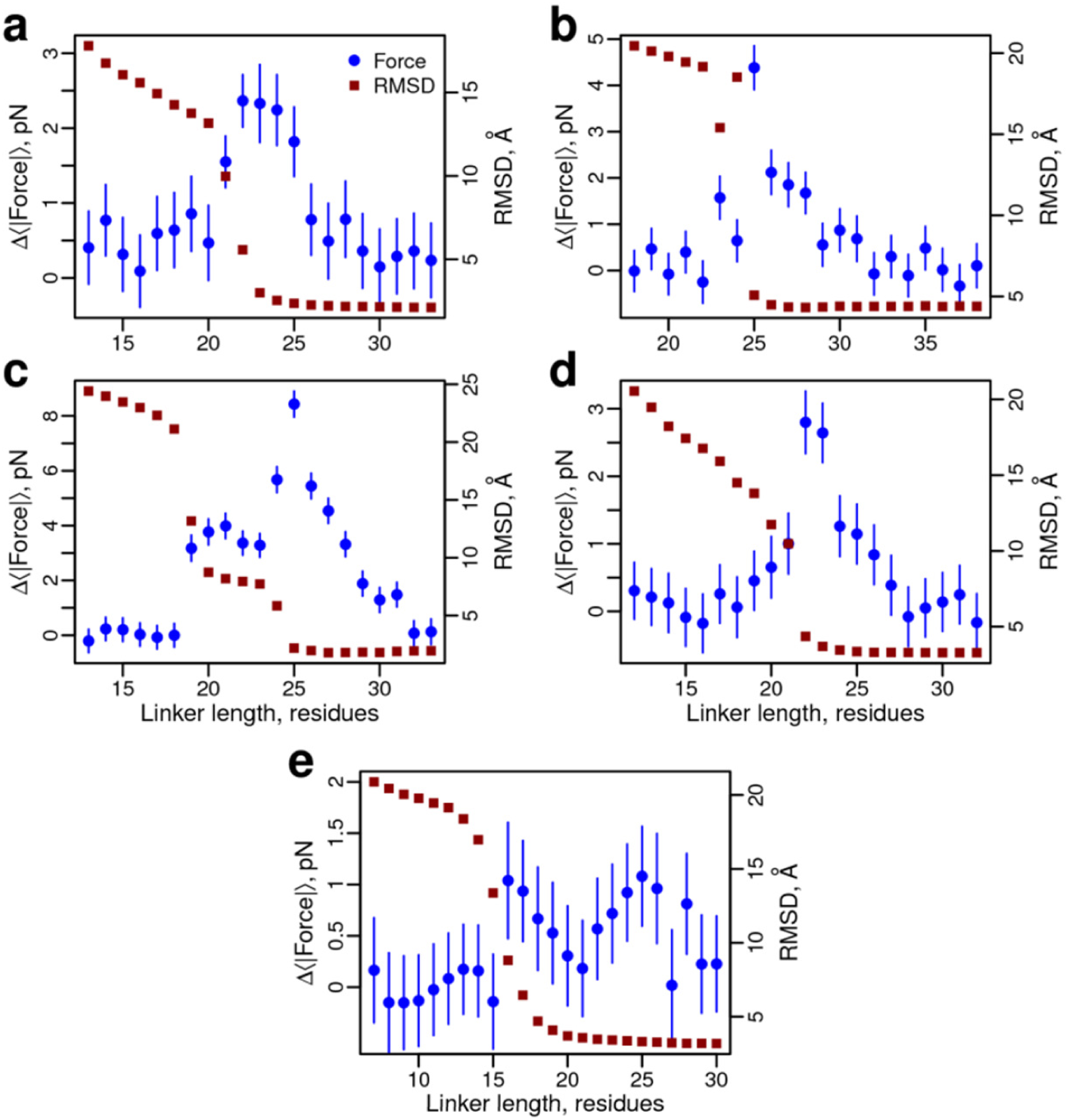
Co-translational folding generates pico-newtons of force at the P-site on translationally arrested ribosomes. Blue circles correspond to the average pulling force at the C-terminal nascent chain residue (Eq. 6) as a function of linker length for the five different proteins shown in Fig. 2: (**a**) 1F0Z, (**b**) 2IST, (**c**) 1P9K, (**d**) 2JSO, and (**e**) 1NY8. Red squares correspond to the average RMSD difference between domain structures sampled in the simulations and the folded crystal structure. Large RMSD values indicate the domain is unfolded, low values indicate it is folded. Note that 1NY8 folds non-cooperatively, and the RMSD fails to report on the docking of the C-terminus beginning at a length of 25 residues, which is seen when using the fraction of native contacts per secondary structural element (SI Appendix, Fig. S1e). All error bars represent 95% confidence intervals about the mean calculated from Block Averaging. Error bars on RMSD data points are smaller than the symbols.

### Non-cooperative folding leads to skewed or bimodal force profiles

Three of the proteins exhibit a single maximum in their force profile (1F0Z, 2IST, 2JSO in Figs. 3a,b and 3d), while protein 1P9K exhibits a shoulder (blue circles, Fig. 3c), and 1NY8 exhibits two maxima (blue circles, Fig. 3e). For the three single-peak proteins we observe they exhibit a cooperative, all-or-none folding transition in their RMSD profiles (*i.e.*, a single inflection point, Figs. 3a,b,d), indicating that they do not populate any co-translational folding intermediates, while 1P9K clearly exhibits non-cooperative folding as indicated by two inflection points in its RMSD profile (red squares, Fig. 3c). 1NY8 also exhibits two force peaks, but has only one visible inflection point in its RMSD profile (red squares, Fig. 3e). As we show below, projecting the simulation trajectories of 1NY8 along other order parameters reveals it folds non-cooperatively.

To determine the origin of the shape of these simulated pulling force profiles we analyzed for each protein the fraction of native contacts of each secondary structural element during synthesis. Consistent with the RMSD results, we find that there is cooperative formation of tertiary structure for proteins 1F0Z, 2IST, and 2JSO (*i.e*., all of the structural elements transition to the native fold at the same linker length), and partially folded intermediates are populated by 1P9K and 1NY8 (SI Appendix, Fig. S1). With this analysis we are able to identify that the folding intermediate of 1P9K consists of tertiary interactions between the three most N-terminal secondary structural elements (two α-helices and a β-strand), while the second transition consists of the docking of the most C-terminal β-strand to form the folded state (SI Appendix, Fig. S1c). The partially folded intermediate of 1NY8 is composed of the six most N-terminal secondary structural elements (three α-helices and three β-strands in SI Appendix Fig. S1e), and the second transition, at a linker length of 25 residues, involves the docking of the last C-terminal α-helix to form the folded state. We emphasize that while the RMSD metric does not reflect this second transition (Fig. 3e), it is revealed using the fraction of native contacts formed by each secondary structural element as an order parameter (SI Appendix, Fig. S1e). In each case, these subdomain folding events correlate with an increase in force, indicating that partially folded intermediates can form and give rise to bimodal and asymmetric force profiles.

These results demonstrate that, for these models, folding generates pulling forces on the piconewton scale 16 to 26 residues after synthesis of the most C-terminal structured residue in the domain, and that formation of folding intermediates can also generate a force.

As a technical aside, because we are reporting the magnitude of the force (Eq. 6), it is possible that we are observing a compressive (pushing) rather than tensile (pulling) force. To confirm it is a tensile force, we calculated the average force vector at the C-terminus of protein 2JSO at linker length 22, which yields a force vector of [33.2, −4.4, −13.2] pN. The angle formed between this vector and the long exit tunnel axis (which lies along the positive X-axis) is 22°, demonstrating that this is a pulling force that is almost parallel with the tunnel and points towards the exit tunnel opening.

Protein folding *in vitro* is influenced by the stability of the domain, its topology, and its size^23,24^. *In vivo*, the speed of protein synthesis can also affect the co-translational folding process^8,10,25–28^. Therefore, we next sought to understand if these factors also influence force generation during co-translational folding.

### Domain stability influences force generation

To study the influence of native state stability we altered the strength of the non-bonded interactions between residues that form contacts in the native state. Greater strength (*i.e.*, a larger Lennard-Jones well-depth, Eq. 3) results in a more stable folded state. This is analogous to experiments in which protein stability is modulated by changing solvent quality using denaturants and osmolytes. For each protein we created at least eight different native stabilities for the protein off the ribosome, ranging between −1 and −8 kcal/mol at 310 K. We then ran arrested ribosome simulations for each stability at the linker length that produced the largest force in the previous simulations (Fig. 3). Protein 1NY8 had a force profile with two maxima that are statistically indistinguishable, therefore we ran simulations at both these linker lengths. We find that for all proteins the pulling force increases with increasing native-state stability, until it plateaus, where further increases in stability do not result in larger pulling forces at that linker length (orange circles in Fig. 4).

**Figure 4.**
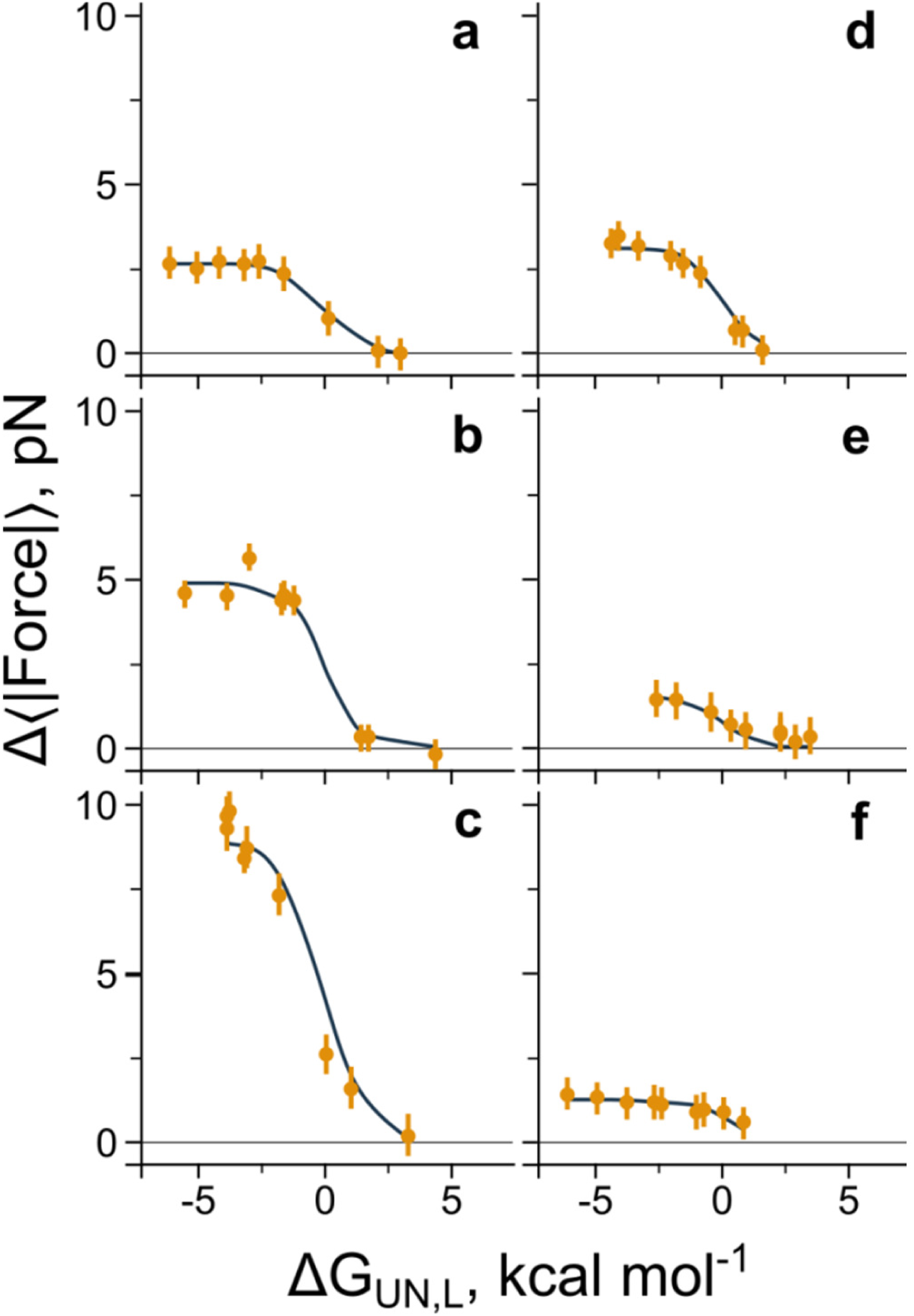
A statistical mechanical model describes how the pulling force changes with native-state stability at a fixed linker length. Pulling force (Eq. 6) versus native state stability for each of the five domains is plotted at a fixed linker length. Panels **a** through **f** correspond, respectively, to proteins 1F0Z, 2IST, 1P9K, 2JSO, 1NY8 and 1NY8 at linker lengths of 23, 25, 25, 22, 16 and 25 residues, respectively. Results from the arrested-ribosome simulations are shown as circles, while the fit to the statistical mechanical model (Eq. 1) is shown as a solid line. Error bars represent 95% confidence intervals about the mean calculated from Block Averaging.

To understand how the pulling force at a fixed linker length changes with native state stability we created a statistical mechanical model in which each domain can exist in one of two states—the native or unfolded state—and that each state generates a characteristic pulling force that is independent of the native-state stability. With these assumptions it can be shown that pulling force is (see SI Appendix)

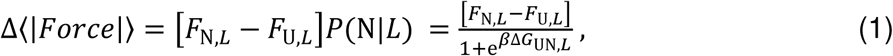

where *F*_N,*L*_ and *F*_U,*L*_ are the characteristic pulling forces of the native and unfolded states respectively at linker length *L, P*(N|*L*) is the probability the domain is in the folded state at linker length *L*, ΔG_UN,*L*_ is the free energy difference between the native and unfolded states at linker length *L*, and 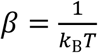, where *k*_B_ is the Boltzmann constant and *T* is the temperature. Using [*F*_N,*L*_ − *F*_U,*L*_] as a fitting parameter, we see this equation reasonably describes how the pulling force changes with stability at a given linker length (solid lines in Fig. 4). (We also provide a generalization of Eq. 1 that describes the folding force when an arbitrary number of intermediates are populated (Eq. S10).) In the limit of an unstable native state, *i.e.*, Δ*G*_UN,*L*_ → ∞, the folded population goes to zero, P(N|*L*) = 0, and therefore Δ〈|*Force*|〉 = 0. At the midpoint of stability P(N|*L*) = 0.5 and hence Δ〈|*Force*|〉 is half its maximal value. And finally when the folded state is very stable, *i.e.*, ΔG_UN,*L*_ → −∞, P(N|*L*) ≈ 1, and the force reaches an effectively constant value of Δ〈|*Force*|〉 = [*F*_N,*L*_ − *F*_U,*L*_].

Thus, a domain’s native state stability influences the magnitude of the pulling force at a fixed nascent chain length. Greater stability leads to greater pulling forces up to a point, beyond which further increases in stability negligibly alter the native state population resulting in no further change in the force at that fixed linker length.

### Domain topology influences force generation

Next, we examined whether domain topology affects force generation. To isolate this effect, we need to control for domain size and stability. Therefore, we took the approach of rewiring the domains, *i.e.* reconnecting the loops between the secondary structural elements of the domain^29^, and adjusted the strength of the non-bonded interactions in the simulation force field so as to maintain iso-stability between the original and rewired forms (see SI Appendix). Domain size is maintained with this procedure as the total number of residues composing the domain stayed constant before and after rewiring; the ordering of the secondary structural elements is the only change. There are many possible ways to rewire the loops, and simulations on the ribosome are computationally expensive. Therefore, we first focused on rewiring one domain based on the hypothesis described below.

Protein 1P9K exhibits the largest folding force (Fig. 3c) and is the only domain in our set of proteins to have a β-hairpin (*i.e.*, two anti-parallel β-strands that are connected by a loop and form tertiary structure) at the C-terminus of the domain (Fig. 5a,b). This β-hairpin segment is the last portion of the domain to emerge from the exit tunnel during synthesis. We analyzed 1P9K’s co-translational folding pathways and found that as the β-hairpin folds (blue circles, SI Appendix Fig. S2a) the more N-terminal β-strand flips back towards the ribosome surface (black squares, SI Appendix Fig. S2b), along with the rest of the domain to form contacts with the more C-terminal β-strand. This process pushes the domain against the ribosome surface, as evidenced by the increased number of contacts between the domain and ribosome (red squares, SI Appendix Fig. S2a), which we hypothesized caused this large pulling force. To test this hypothesis we rewired the loops to move the C-terminal β-hairpin to the center of the domain (Fig. 5a), while maintaining the same folded-state stability off the ribosome. The average RMSD between the secondary structural elements in the original and rewired domain is 1.6 Å at 310 K, thus the overall shape of the protein is not perturbed by rewiring. We find that moving the β-hairpin in this way causes the maximum force to decrease from 8.5 pN to 3.8 pN (Fig. 5c), thus supporting our hypothesis. As a further test of this hypothesis we took protein 1NY8, which has a maximum force of 1.1 pN and no C-terminal β-hairpin, and rewired its loops to create a C-terminal hairpin (Figs. 5d,e) while maintaining its stability. We find the maximum force increases to 8.9 pN (Fig. 5f), again supporting our hypothesis.

**Figure 5.**
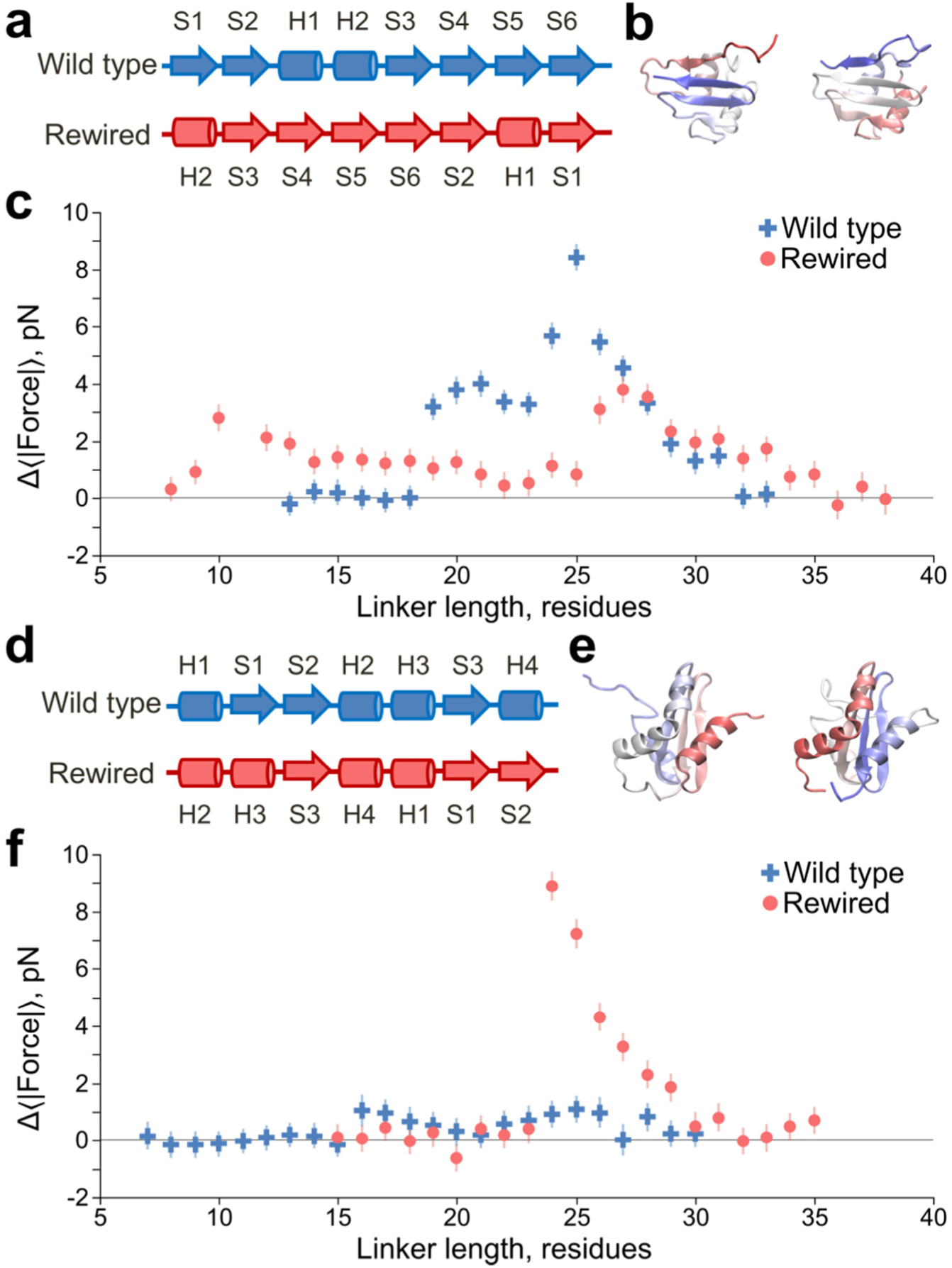
Topology influences the pulling force, with C-terminal β-hairpins giving rise to large forces. **(a)** Linear secondary structure representation, from N-to C-terminus, of the domain 1P9K. Arrows indicate β-strands, cylinders indicate α-helices. The original wild-type structure is shown at the top, while the rewired mutant is shown below. The rewired mutant maintains the same secondary structure labels (“H1”, “S1”, *etc.*) as the wild-type to illustrate how their relative positioning has changed along the primary structure due to rewiring. **(b)** Crystal structures of the wild-type (left) and rewired (right) domains, with the N-terminii in red and the C-terminii in blue. **(c)** Pulling force (Eq. 6) versus linker length for the wild-type (blue circles) and rewired mutant (red squares) domains at 310 K. Moving the C-terminal β-hairpin (formed by strands S5-S6) to the middle of the domain generates a lower maximum force in the rewired mutant. **(d)** Same as (a) except for protein 1NY8, which in the wild-type has no C-terminal β-hairpin, but in the rewired mutant has one formed by strands S1-S2. **(e)** Same as (b) except for protein 1NY8. **(f)** Same as (c) but for protein 1NY8; wild-type results are blue circles, mutant results are red squares. Error bars represent 95% confidence intervals about the mean calculated from Block Averaging.

These results demonstrate that topology can influence the pulling force, and that the presence of C-terminal β-hairpins can generate relatively large forces.

### Domain size has no apparent effect

Polymer theory models and experimental observation demonstrate that an entropic pulling force can be generated when an inert sphere is attached to the end of an unbranched polymer that is, itself, covalently attached to a surface^30,31^. This pulling force increases as the radius of the bead increases until its size becomes comparable to the polymer’s contour length, after which this force approaches a constant value^30^. We examined whether a similar size-dependent force might arise due to the size of the folded domain (the sphere) attached to the linker (the polymer) that is attached in turn to the ribosome (the surface). To do this we controlled for stability effects by tuning all five domains to a native state stability of −4.5 kcal/mol off the ribosome and determined the maximum pulling force generated during co-translational folding, again on arrested ribosomes. No correlation between force and domain size is found when domain size is defined as the number of residues in the domain (Fig. 6) nor by its radius of gyration (SI Appendix, Fig. S3), suggesting domain size has no effect. We emphasize that this result does not definitively rule out an effect from domain size. We have seen that domain topology can have significant effects on the magnitude of the pulling force. Since we did not control for differences in topology between domains, any contribution from domain size has the potential to be obscured. Additionally, the polymer model^30^ predicts that for the domain sizes used in this study there would only be a 0.25 pN force difference between the smallest and largest domain. A force that again could be obscured by topological differences.

**Figure 6.**
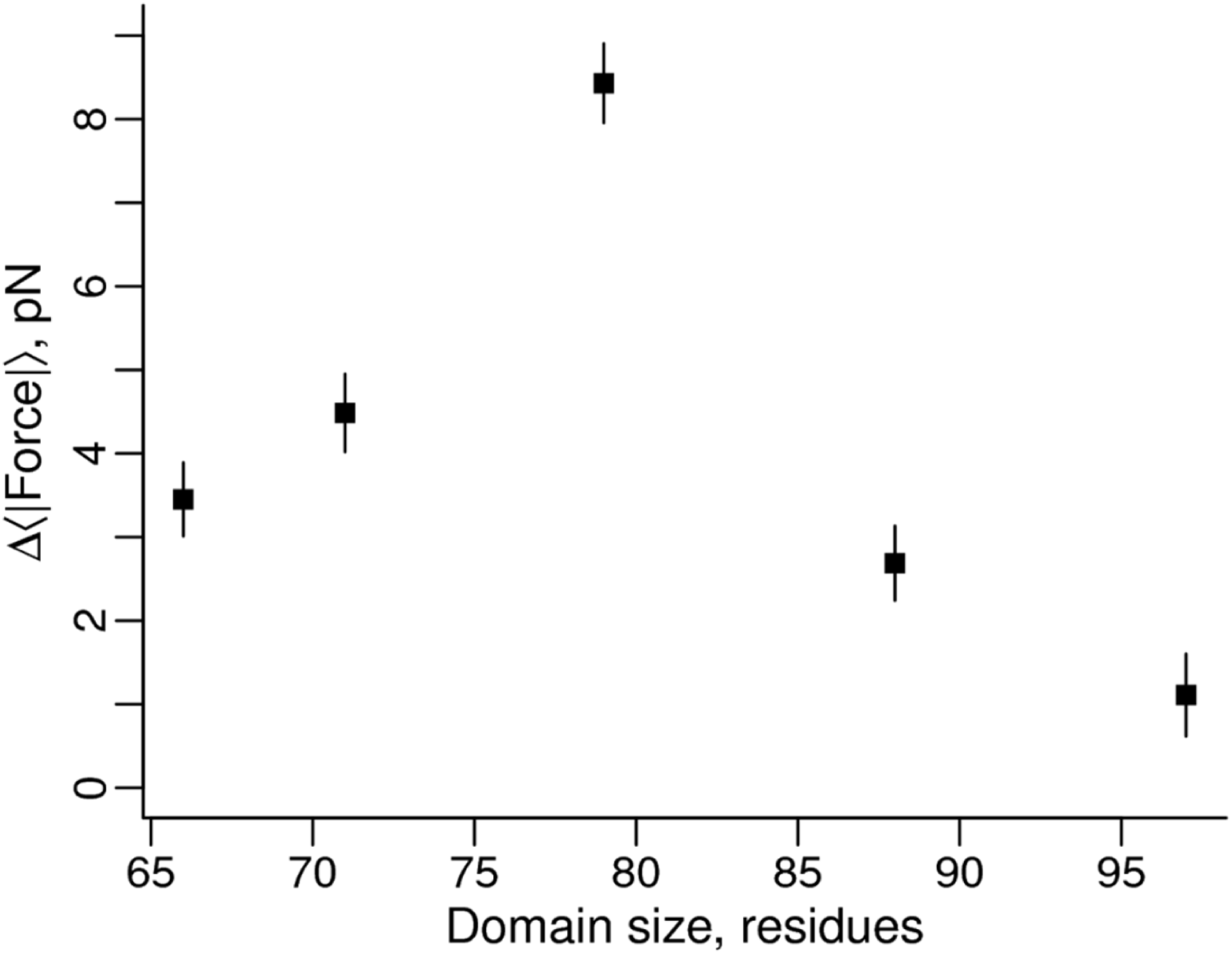
The size of the domain has no apparent effect on the magnitude of the force. Pulling force versus domain size for the five different domains. No statistically significant trend is observed. Error bars represent 95% confidence intervals about the mean calculated from Block Averaging.

### Translation speed influences force generation

The results we have reported thus far are based on arrested ribosome simulations in which the co-translational folding process is at equilibrium at each nascent chain length. Continuous synthesis, being irreversible, is a non-equilibrium process and therefore folding, and hence mechanical forces, have the potential to differ between arrested and continuous translation^25,32^. To test if the folding force differs between these two situations we ran continuous synthesis simulations of proteins 2JSO and 2IST (Fig. 7). We emphasize that while our coarse-grained simulations exhibit accelerated dynamics^33^, we maintain a realistic ratio of time scales between domain folding and amino-acid addition (see Eq. 4), which allows us to estimate that 50 ms of experimental time corresponds to 12.6 ns of simulation time based on the average value of α from SI Appendix Table S1 (see also Eq. 5). For 2JSO, the force profile obtained from continuous synthesis simulations is consistent with the force profile from the arrested ribosome-nascent chain simulations (Fig. 7a). 2IST’s force profile, however, generated no force during continuous synthesis (red squares, Fig. 7b). When continuous translation was further slowed down by using the median value of α, the pulling force remained lower than that found on the arrested ribosome (gray triangles, Fig. 7b). Thus, translation speed can decrease or abolish the pulling force generated by proteins on arrested ribosomes.

**Figure 7.**
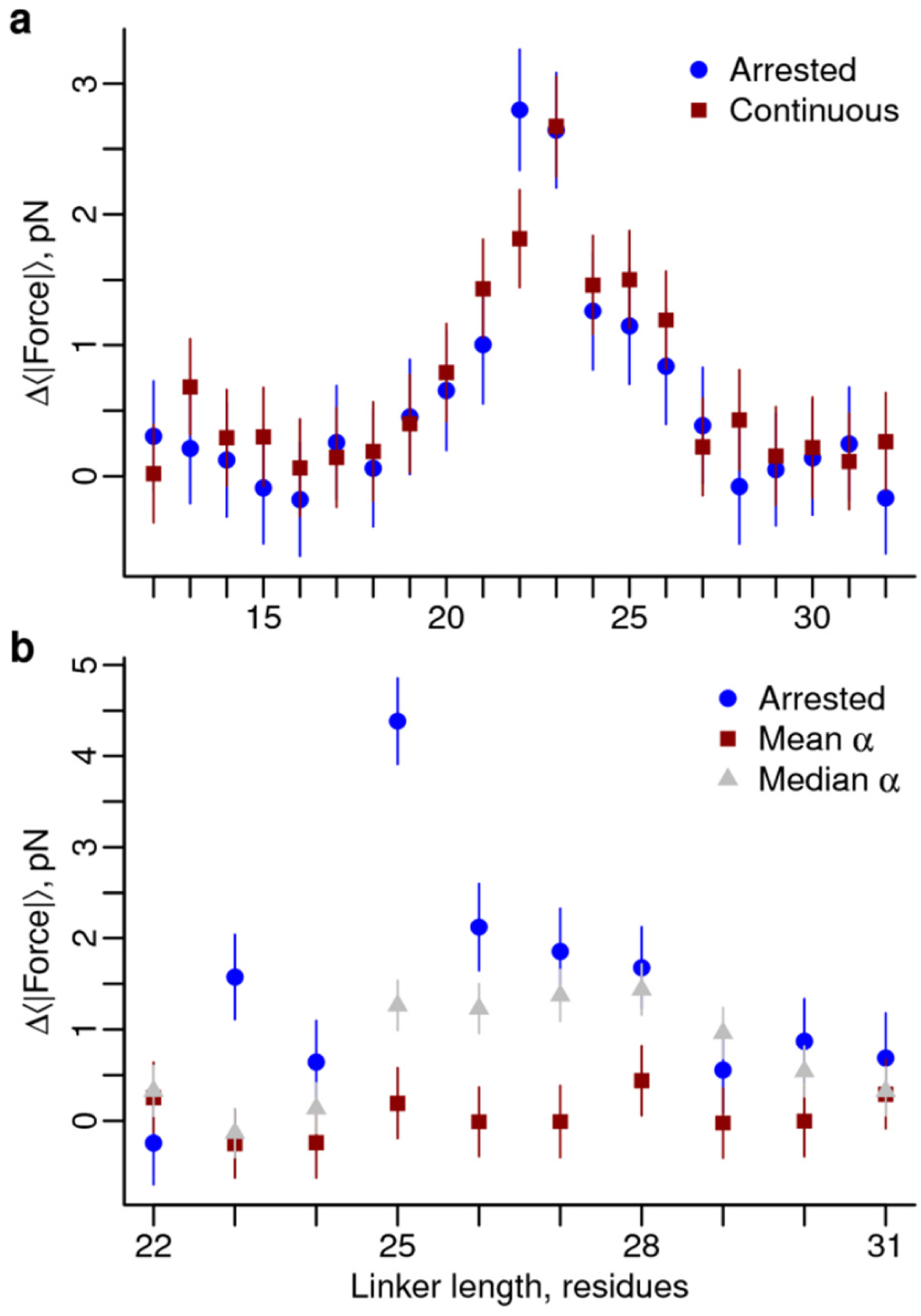
Pulling forces generated on arrested ribosomes can differ from those undergoing active translation. **(a)** Pulling force versus linker length for protein 2JSO on arrested ribosomes (blue circles) and undergoing continuous synthesis (red squares). **(b)** Same as (a) except for protein 2IST and two translation rates, the mean α (red squares) and the median α (gray triangles). The force profiles arising from arrested and continuous synthesis are in good agreement for 2JSO, while for 2IST the pulling force decreases during continuous synthesis, and the extent of change depends on the exact translation rates. Error bars represent 95% confidence intervals about the mean calculated from Block Averaging (arrested simulations) or Bootstrapping (continuous simulations). Using a Wilcoxon signed-rank test to compare the rates in this folding region, p<0.05 between the arrested and median α, and p<0.001 between the mean α and median α, and between the arrested and mean α.

During continuous translation the co-translational folding of a domain occurs at longer nascent chain lengths when the mean folding time of the domain (*τ*_N_) is longer than the timescale at which amino acids are added to the elongating chain (*τ*_A_) by the ribosome^25^. We hypothesized that such a delay in folding was causing 2IST’s pulling force to decrease during continuous synthesis. To test this, we calculated the mean folding time of the 2IST and 2JSO proteins at a range of linker lengths that bracket the midpoint of folding in the arrested ribosome simulations. We find that while 2IST folds 2 times faster than 2JSO off the ribosome, on the ribosome 2JSO’s ratio of *τ*_N_/*τ*_A_ is less than 1 at a majority of linker lengths (*i.e*., folding is faster than amino acid addition) while 2IST’s ratio is greater than 1 up to seven residues beyond the midpoint length of folding (Fig. 8a). This has the effect of shifting 2IST’s midpoint of folding to six residues later during continuous synthesis as compared to on an arrested ribosome (Fig. 8b). This delay in folding means that when 2IST finally folds during continuous synthesis it is no longer pushing against the ribosome surface, and hence cannot generate a pulling force. This interpretation is supported from the simulations: at a linker length of 24 residues the folded state of 2IST is in contact with the ribosome surface 100% of the time, while at linker length 30 the folded state is only in contact 40% of the time.

**Figure 8.**
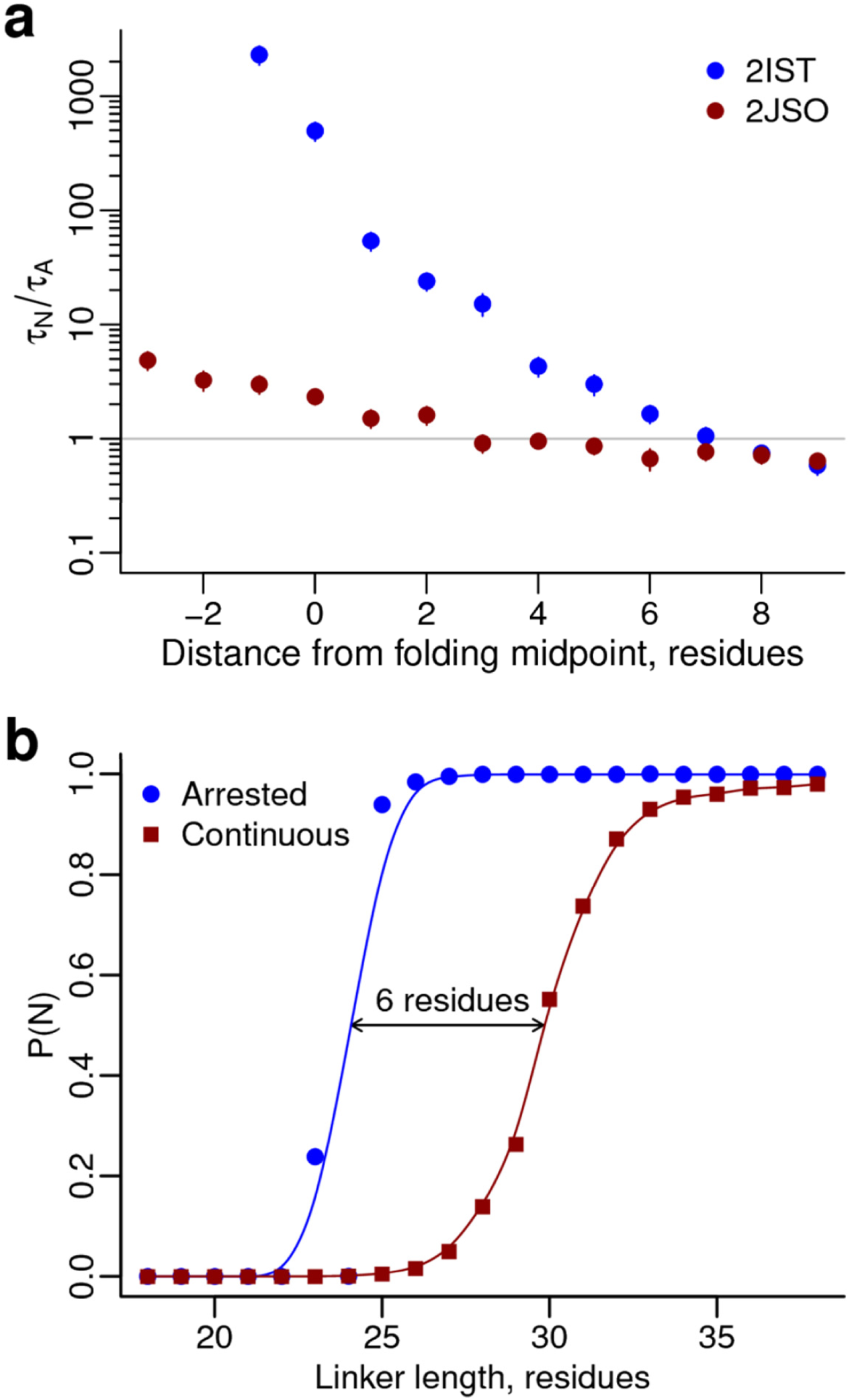
A delay in co-translational folding due to continuous synthesis causes 2IST’s pulling force to vanish. **(a)** The ratio of folding time to amino acid addition time versus linker length for proteins 2IST (blue circles) and 2JSO (red circles) as calculated from temperature quenching simulations on arrested ribosomes. While 2IST folds faster in bulk solution, its folding rate slows at short linker lengths on the ribosome. Error bars represent 95% confidence intervals about the mean, calculated from Bootstrapping. **(b)** Probability of folding versus linker length for protein 2IST on arrested ribosomes (blue circles) and undergoing continuous synthesis (red squares). The longer folding time for 2IST causes the protein to fold six residues later during continuous synthesis than on an arrested ribosome.

Thus, translation speed can influence force generation when the time scale of folding is longer than the amino acid addition timescale, which shifts the folding process from a quasi-equilibrium to a non-equilibrium regime^32^. As little as a six-residue delay in folding can diminish or abolish force generation.

### Statistical mechanical model explains trends in force profiles

We demonstrated that a statistical mechanical model can describe how the pulling force changes with native-state stability at a given nascent chain length (Fig. 4). If we understand force generation during synthesis we should be able to create a model that predicts the pulling force at all linker lengths (*L*) during arrested or continuous synthesis. To create this model (see SI Appendix for derivation) we calculate the intrinsic difference in force between the native (N) and unfolded (U) states [*F*_N_ - *F*_U_], and multiply it by the probability the domain is folded at a given length, P(N|*L*), and by the conditional probability that, given the domain is folded and touching the ribosome (*T*), it does not have slack in the linker,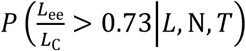. Based on a polymer physics model of a Worm-like Chain that exhibits a cross-over to a high-force regime in its force versus extension curve (brown arrow in Fig. 9a)^34^, we define slack as being present if the linker’s end-to-end distance, *L*_ee_, is less than 73% of its contour length, *L*_C_ (Fig. 9a). When [*F*_N_ - *F*_U_] is used as a fitting parameter, the product 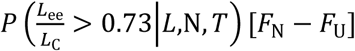 is consistent with the raw [*F*_N_ - *F*_U_] values calculated directly from the simulations. P(N|*L*) is related to the folding (*k*_N_(*L*)) and unfolding (*k*_U_(*L*)) rates of the domain, and the codon translation rates (ω _A_(*L* + 1)) at a given length by the function 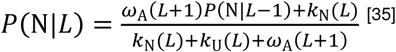. The resulting equation is

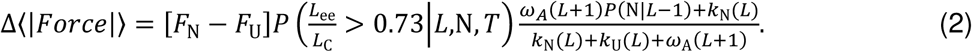

**Figure 9.**
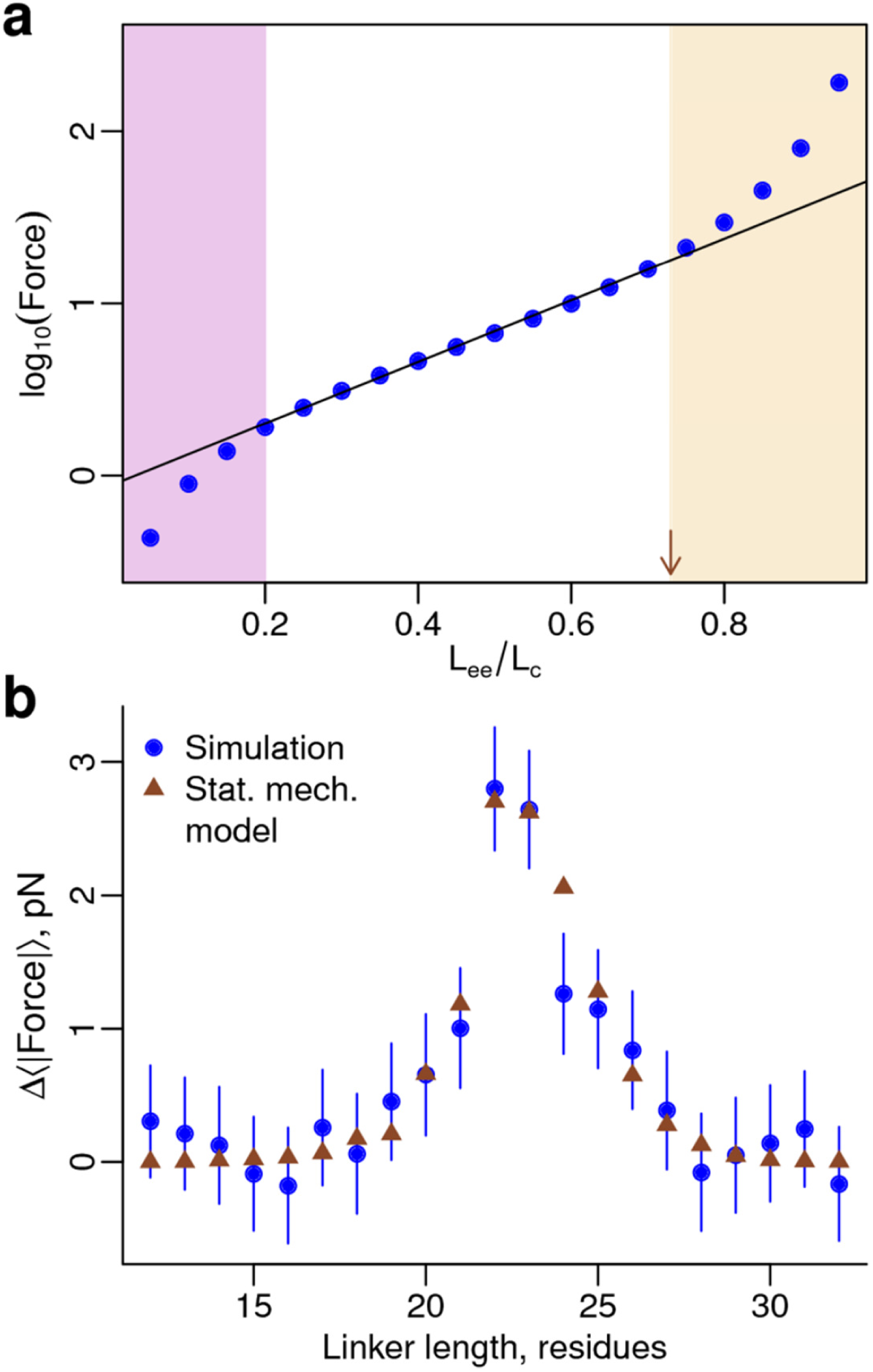
A statistical mechanical model describes how the pulling force changes with linker length on arrested ribosomes. **(a)** A force versus extension curve for a Worm-like Chain polymer with a persistence length of 9 Å, contour length (L_c_) of 84 Å and confined to an inert cylinder with a 15 Å diameter at 310 K^34^. The abscissa plots the end-to-end extension of the polymer (L_ee_) divided by its contour length. Low, intermediate, and high force regimes are present, with the line of best fit to the intermediate regime demarcating it from the high force regime at L_ee_/L_C_ = 0.73, red arrow. As noted in the Methods, this value was used as a threshold. (**b**) The statistical mechanical model (Eq. 2) accurately describes the force versus linker length results from the simulations of protein 2JSO. Error bars represent 95% confidence intervals about the mean calculated from Block Averaging.

We note that in the limit of infinitely slow translation (*ω*_*A*_(*L* + 1) = 0, *i.e.*, an arrested ribosome), 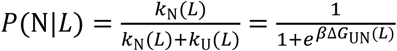, where Δ*G*_UN_(*L*) is the free energy as a function of linker length, and we get back that the force is a function of the native-state free energy at each length.

We first tested the accuracy of this equation by comparing its predicted force profile to that from the arrested ribosome simulations of protein 2JSO. Setting ω _A_(*L* + 1) = 0 and plugging in *F*_N_, *F*_U_, Δ*G*_UN_(*L*), and 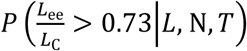, which were determined from the simulations (see SI Appendix), we find that Eq. 2 reasonably describes 2JSO’s force profile (Fig. 9b).

Next, we tested how this equation performs in describing the continuous synthesis force profile from 2IST. Using the same codon translation rates as in the simulation, we find Eq. 2 correctly predicts 2IST will not produce a force at these rates (black squares, Fig. 10), and will produce a force when translation is very slow compared to folding (red triangles, Fig. 10). We illustrate in Fig. 10 (blue circles) that Eq. 2 can also predict force profiles at intermediate translation speeds.

**Figure 10.**
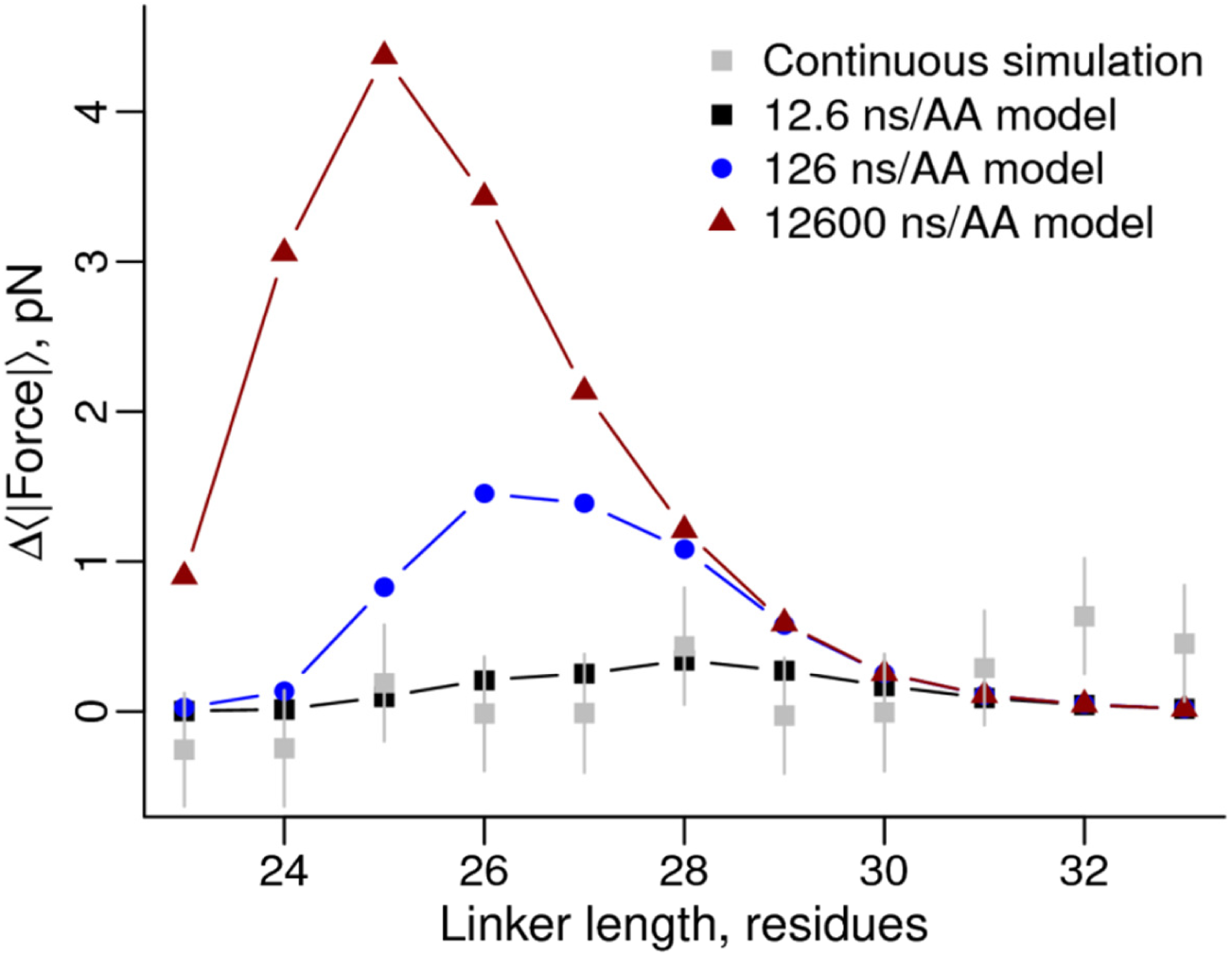
A statistical mechanical model describes the influence of translation speed on force. We calculated folding probabilities for protein 2IST at a range of linker lengths for various amino acid addition rates and used the statistical mechanical model Eq. 2, to estimate the pulling force at each length and speed. The black squares represent the rate used in the continuous synthesis simulations, while the blue circles and red triangles represent one and three orders of magnitude faster, respectively. Error bars represent 95% confidence intervals about the mean, calculated from Bootstrapping.

Thus, Eq. 2 is a non-equilibrium statistical mechanical model that can describe co-translational folding force generation on both arrested and continuously translating ribosomes.

## Discussion

Tensile forces acting on the nascent protein can alter the speed of translation elongation in a form of mechanical allosteric communication that links what happens outside the exit tunnel to the ribosome’s catalytic core^1–7^. It has been found that protein folding on translationally arrested ribosomes can generate a pulling force that is transmitted through the nascent protein backbone to the peptidyl transferase center (PTC), which can alter peptide bond formation^1,4,7^. Specifically, a previous study used Quantum Mechanics/Molecular Mechanics simulations to show that pulling forces applied to the PTC can decrease the free energy of the transition state barrier to peptide bond formation^7^. The results from this coarse-grained study quantify this novel form of allosteric communication within the ribosome-nascent-chain complex, identify the factors influencing the strength of this allosteric signal, and present statistical mechanical models to predict these forces. This tensile-force feedback mechanism may potentially be a common occurrence during protein synthesis, as it is estimated that one-third of the *E. coli* proteome co-translationally folds^25^, and these results lay the ground work for future research exploring that possibility. The domain properties and series of molecular events that give rise to this force and the influence of continuous translation were previously unknown. Understanding the impact of these factors provides insight into this source of mechanochemistry on the ribosome and how it could influence translation speed, and opens up avenues to understanding the evolutionary pressures that have shaped properties of domains.

In this study we examined four different factors that we hypothesized had the potential to influence mechanical force generation on the ribosome. Utilizing a coarse-grained simulation model and statistical mechanics, we demonstrated that a domain’s stability and topology, as well as translation speed, determine co-translational folding force generation on the ribosome. A number of theoretical and experimental studies have demonstrated that the pulling force a polymer experiences as it is extruded through a pore is proportional to the change in free energy of the polymer during translocation^36–38^. This is broadly consistent with our finding that the free energy of folding influences the pulling force.

Topology is another factor that governs the pulling force. We found that inserting and removing C-terminal β-hairpins increased and decreased the pulling force, respectively. This is because to form a C-terminal β-hairpin, the rest of the domain (if it is already folded) must flip back toward the ribosome surface and press against it. To our knowledge this is the first time this topological effect has been observed in either simulations or experiments, and suggests the possibility that the presence of C-terminal β-hairpins in domains may have been naturally selected for in domain structures to modulate translation speed. Our study of five α/β proteins is not comprehensive and therefore other topological features may also have systematic effects on force. For example, the presence of C-terminal α-hairpins might have a similar effect since a similar series of structural transitions might occur (SI Appendix, Fig. S2).

Translation speed was found to have no effect on the pulling force when a domain’s folding time was equal to or less than the codon translation time, but diminished the force when folding was slower than the elongation time. It was previously shown that slow-folding domains fold at longer nascent chain lengths during continuous synthesis compared to the folding of those same domains on arrested ribosomes. And that for 20% of cytosolic *E. coli* proteins, translational kinetics can switch their domains from folding co-translationally to post-translationally^25^. We found that protein 2IST’s slow folding time caused it to fold at longer nascent chain lengths, and that when it folded it was no longer near the ribosome surface to press against it and generate a pulling force. This suggests two things - for experiments, some of the force measurements on arrested ribosomes may not be relevant to continuous synthesis, especially if the domains fold slower than the codon translation times (*i.e.*, slower than about 200 ms^39^). And that forces may generally tend to be lower during continuous synthesis. For example, 2IST’s force completely disappears during continuous synthesis simulations.

Due to limitations in experimental spatial resolution the structural basis for force generation is unknown, although a number of hypotheses have been stated in the literature^1,2,35^. Our simulations provide insight into the structural changes that occur concomitant with force generation, and our statistical mechanical modeling reveals the essential features. For example, if all we consider is the folding status of the domain during synthesis, then we cannot accurately model the force profiles seen in the simulations (blue circles, SI Appendix Fig. S4). If we then speculate that we need to know not just whether the domain is folded but also when it is touching and pressing against the surface of the ribosome to generate a force, we still cannot accurately describe the force profile (orange diamonds, SI Appendix Fig. S4). The reason for this is that there are many structures where the domain is folded and touching the surface but the linker has slack in it. It is only when we require that the domain be folded, pressing against the surface, and have little slack in the linker that we see good agreement between the predicted force profile and the measured force profile (black squares, SI Appendix Fig. S4). Thus, at the molecular level, forces are generated just as a domain emerges from the exit tunnel, begins to fold, pushes against the ribosome surface, and there is little slack in the linker. All three of these structural features of the domain and linker are essential components of force generation.

We observed a maximum pulling force of between 1 and 9 pN. The only experimentally estimated pulling force was slightly higher at 12 pN for the protein Top7^1^. Several features of the Top7 protein help to explain why its force is larger. The Top7 domain is an engineered domain that is highly stable for its small size—the free energy of folding is −13.2 kcal/mol, which is more than twice the free energy of folding of our most stable domain^40^. Additionally, the Top7 protein contains a β-hairpin at its C-terminus^40^, just like our two highest force-producing constructs. Thus, a larger force should be expected for Top7 compared to the proteins we studied.

Two unexpected results from this study are the lack of any effect from domain size and the differing effect of the ribosome on folding rates. Larger domains were predicted to exhibit a greater pulling force due to the larger increase in entropy they can achieve by moving away from the ribosome surface. We observe no correlation, however, between the maximum force and domain size when we control for differences in domain stability. We emphasize that this result does not definitively rule out an effect from domain size. We have seen that domain topology can have significant effects on the magnitude of the pulling force. Since we did not control for topology in this comparison (because it is not possible to isotropically change a domain’s size), any contribution from domain size has the potential to be obscured by topological effects, especially in light of the force differences predicted by the polymer theory model^30^. Less germane to the purpose of this study, we observed, to our knowledge, the first example of a change in ordering of fast- and slow-folding proteins when they fold co-translationally. Specifically, we observe that off the ribosome protein 2IST folds 2.2 fold faster than protein 2JSO. But on the ribosome, protein 2JSO can fold 210-fold faster than protein 2IST (compare points at an abscissa value of 0 in Fig. 8a). This indicates differences in native state topologies can play an important role in folding kinetics on the ribosome.

A key technical point we wish to convey is the challenge of acquiring equilibrium results from the arrested ribosome simulations even for these relatively small domains. If the simulations are not at equilibrium, the calculated force can be inaccurately very high or low. A key test for equilibrium sampling in the replica exchange (REX) simulations used here is observing a constant number of transitions from the high to low temperature replicas in different portions of the simulation. For example, for protein 1F0Z at a linker length of 24 residues, we observe a constant number of transitions in the various deciles of the replica exchange simulation (SI Appendix, Fig. S5b). If there was non-equilibrium sampling this quantity would exhibit a systematic drift with time. In our view, to help ensure accurate results, such a demonstration of equilibration should be required of any REX simulations from which forces or other thermodynamic averages are calculated.

We have used simulations and created statistical mechanical models to identify trends in the pulling force arising from various factors. Our results concerning the influence of domain stability, topology, and translation rate could be tested experimentally using expression assays comparing the amount of full-length protein produced to the amount of truncated protein produced due to translation stalling by SecM. To test stability, domains could be expressed in solvents of varying solvent quality^41^, or point mutations could be introduced^42^. Translation speed effects might be tested by introducing synonymous codon mutations in the mRNA^43^, and altering the topology using circular permutation is a common technique in protein engineering^29,44^.

In summary, we have identified the factors determining mechanical force generation on the ribosome due to co-translational protein folding, created statistical mechanical models that describe how force profiles depend on these factors, and described the series of molecular events giving rise to this force. In future studies it would be interesting to probe proteome-wide structural domain properties and transcriptome-wide codon usage bias to look for signatures of evolutionary selection acting on these properties, and the biological benefit of this form of translation speed modulation through mechanochemistry.

## Methods

### Ribosome and nascent chain models

Five cytosolic *E. coli* proteins were randomly selected from the Protein Data Bank (PDB) with the requirements that they be of the α/β structural class, bind no ligands, have no post-translational modifications, and are between 50 and 100 residues in length. This resulted in four single-domain proteins (PDB IDs 1F0Z, 1P9K, 1NY8, 2JSO), while the fifth was a two-domain protein, but only the N-terminal domain was studied here (PDB 2IST, residues 2-72).

Coarse-grained structure-based models were created for each domain as previously described^45^, where each amino acid is represented as a single interaction site centered on its α-carbon and assigned a +/-1 or 0 e^-^ charge based on the overall charge of the amino acid at pH 7. Counter ion screening is represented implicitly using Debye-Hückel theory with a screening length of 10 Å. The starting structure for coarse-graining the 50S subunit of the *E. coli* ribosome was PDB ID 3UOS^46^. Ribosomal proteins were coarse-grained in the same manner as the nascent proteins. Ribosomal RNA was represented as three or four interactions sites per nucleotide with a −1 charge assigned to the phosphate center, and for efficiency, the bulk of the ribosome is fixed in space and interacts with the nascent chain only via excluded volume and electrostatic interactions. Residues 42 to 59 of the ribosomal L24 protein were not fixed and allowed to fluctuate since these residues sit at the end of the exit tunnel and can affect co-translational folding. The ribosome was trimmed in an umbrella shape, with the PTC at the base of the narrow section (SI Appendix, Fig. S6). Ribosomal beads within 10 Å of the walls of the exit tunnel and a large portion of the outer ribosomal surface surrounding the exit tunnel were included in the final structure, resulting in a coarse-grained ribosome structure consisting of 6,612 interaction sites.

All simulations were performed using CHARMM version c35b5^47^, which expresses the potential energy of a system as

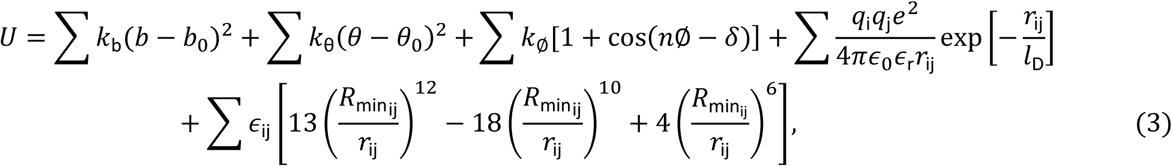

where, from left to right, the summations are over bond, bond-angle, dihedral, electrostatic, and van der Waals energy terms. Parameters for each energy term in Eq. 3 were taken from ref. 45. Note that the parameters for the first four terms are fully transferable between proteins, while the last term is structure-based and hence not transferable. For all simulations the system was propagated in time using Langevin dynamics with a friction coefficient of 0.05 ps^-1^ and an integration timestep of 0.015 ps. To tune the native-state stability of each domain to the value predicted by the PREFUR method^21^, *∈*_ij_ in Eq. 3 was multiplied by a number η. A range of η values were tested for which the thermodynamic properties of each domain were calculated using REX simulations^48^. At least 50,000 exchanges were attempted in these simulations that had at least eight temperature windows, with the first 10,000 exchanges discarded to allow for equilibration. Free energies of stability were calculated using the Weighted Histogram Analysis Method (WHAM)^49^, and η was adjusted until the native-state stability in the model matched the desired stability.

### Arrested ribosome simulations

After each protein domain was tuned to its target stability, an unstructured peptide linker between 1 and 40 residues in length was covalently attached to the C-terminus of the domain (SI Appendix, Table S2)^50^, and the C-terminus of the resulting nascent chain was harmonically restrained with a force of 10 kcal/mol/Å^2^ at the P-site location found in the crystal structure. This restraint was used because it allowed for the fastest equilibration of the force. This value should not affect the value of the pulling force because the pulling force is the difference of the magnitudes of the wild-type and destabilized versions of the domain, removing the influence of the restraint. For each wild-type sequence REX simulations were carried out consisting of between 8 and 14 temperature windows and 300,000 attempted replica exchanges. For the destabilized proteins only 100,000 exchanges were attempted because no phase transition can occur. The first 10,000 exchanges were discarded to permit equilibration. Error bars were computed for REX simulations by breaking each simulation into blocks of 50 exchanges and using the average from each block to compute 95% confidence intervals. 50 exchanges per block were used as it resulted in uncorrelated data^51^.

### Mapping simulation time to experimental time

Coarse-graining and low-friction Langevin dynamics accelerate processes. To map these accelerated dynamics to real-world timescales we simulated the folding of 18 proteins with our coarse-grained model. By comparing the simulated folding time to the experimentally measured value we determined the proportionality constant between the simulation and experimental time scales. Specifically, coarse-grained models, as previously described, were built for six α-helical (SI Appendix, Table S3), six β-sheet (SI Appendix, Table S4), and seven α/β proteins (SI Appendix, Table S5) whose lengths vary from 58 to 155 residues and for which free energy changes upon folding^52,53^ 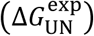 and mean folding times^52,54–58^ 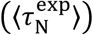 have been experimentally measured. Three to six REX simulations with various η values were carried out for each protein and the free-energy of folding 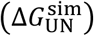 computed for each value of η using the WHAM equations. The fraction of native contacts (*Q*) was used to differentiate folded and unfolded conformations. The *Q* threshold separating the N and U populations at equilibrium, *Q*_eq_, was identified as the value of *Q* at which the cumulative probability of *Q* at the model’s melting temperature (*T*_M_) equals 0.5. The 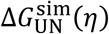 data for each protein were then fit to the equation 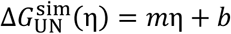 and the η that results in 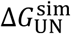 matching the experimental stability, denoted η*, was calculated as 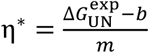, where *m* and *b* are the slope and y-axis intercept, respectively. All values of 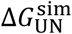 were computed at the temperature at which 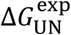 was obtained (*T*_exp_). The η^∗^ values calculated by this method range from 0.998 to 1.628, with an average value of 1.235 (SI Appendix, Table S3-S5).

CG models were constructed for all 19 proteins at their respective η^∗^ values. The mean time of folding for each protein, denoted 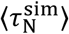, was then determined from temperature quench simulations. Five-hundred independent trajectories were run, with a 1000-K equilibration period of between 20 ns and 2 µs followed by a quench to 310 K lasting between 100 ns to 10 µs. The durations of these two phases of the simulation were chosen such that the length of the 1000 K equilibration was 20% the duration of the 310-K quench. Kinetic *Q* thresholds, denoted *Q*_kin_, were defined for each protein to differentiate the unfolded and folded ensembles during the quench period. This parameter was calculated as the arithmetic mean of *Q*_eq_ and the most probable value of *Q* observed at 310 K (*Q*_310_) in the REX simulation at the value of 1 closest to η^∗^ for a given protein. We required that the folded state remain folded for at least 150 ps before the protein was considered to have folded at the initial frame at which *Q* > *Q*_kin_. The time-dependent survival probability of the unfolded state was calculated as 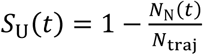, where *N*_N_(*t*) is the number of trajectories that have populated the folded state based on the above kinetic definition of folding at least once by simulation time *t* and *N*_traj_ is the number of statistically independent trajectories included in the analysis. Survival probability curves for all proteins except ABP1 SH3 (PDB ID: 1JO8) were fit to the equation *S*_U_(*t*) = exp(-*k*_N_ ∗ *t*) using Python, where *k*_N_ is the sole fitting parameter, and the mean folding time calculated as 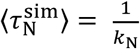. The survival probability distribution for ABP1 SH3 was found to be best fit by the sum of two exponential terms; the fit equation for ABP1 SH3 was modified to the form *S*_U_(*t*) = *f*_1_ exp(-*k*_N,1_*t*) + *f*_2_ exp(-*k*_N,2_*t*) where *f*_1_ + *f*_2_ = 1. Curve fitting revealed that *f*_*1*_ = 0.49, *f*_2_ = 0.51, *k*_1_ = 9.87 × 10^-3^ ns^-1^, and *k*_2_ = 2.18 × 10^-1^ ns^-1^; the value of 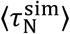 reported in SI Appendix Table S5 for ABP1 SH3 was calculated as 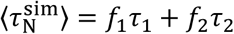 where 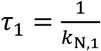 and 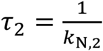. Pearson *R*^2^ values for all fits were ≥0.98. Note that very few trajectories for dihydrofolate reductase (PDB ID: 1RX4) folded during 10 µs of quench and 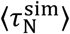 therefore could not be calculated.

We maintain the experimentally observed ratio of a protein’s folding time to the timescale of amino acid addition by requiring that

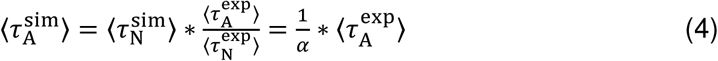

in which we have introduced the parameter 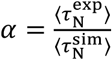 which reports the relative acceleration of protein folding observed in our simulations compared to experiment. The 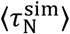, *α*, and other relevant parameters for each of the 18 proteins are reported in SI Appendix Table S5. The mean value of *α* over the 18 proteins in the data set was found to be 〈 *α* 〉 = 3,967,486. The average codon translation rate in *E. coli* increases from 8 to 22 amino acids per second as *E. coli*’s growth rate increases^59^. Therefore, we set 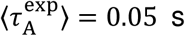 (corresponding to 20 AA/s) yielding,

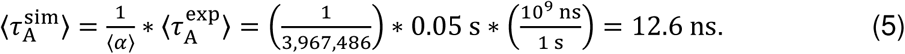

That is, 12.6 ns of simulation time equals 50 ms of experimental time. We emphasize that the acceleration of folding we observe, and therefore this mean *in silico* translation time, is dependent on the specifics of our coarse-grained model (*i.e.*, the force field, frictional coefficient, temperature, model resolution, *etc*.) and is thus not transferrable to other coarse-grained models. On a side note, bacterial translation rates are faster than eukaryotic translation rates due to their faster growth rates. For example, *S. cerevisae* and human cells have translation rates between 5 and 9 amino acids per second^60,61^.

### Continuous synthesis simulations

For the continuous synthesis simulations, 1,100 independent trajectories were run for the wild-type and destabilized versions of proteins 2JSO and 2IST starting from a nascent chain length of 40 residues and ending at 138 and 121 residues, respectively. The same unstructured linker as in the arrested ribosome simulations was added to the C-terminus of the domain and synthesized starting from 89 residues, for 2JSO, and 72 residues, for 2IST, which allowed the domains to fully emerge from the exit tunnel. Residues were added every 12.6 ns on average, with dwell times stochastically sampled from an exponential distribution. Error bars on average properties calculated from these simulation represent 95% confidence intervals computed from bootstrapping 10,000 times.

### Analyses

The *x, y*, and *z* components of the force (*f*_*x*_, *f*_*y*_, *f*_*z*_) from the harmonic restraint applied to the C-terminal bead of the nascent chain was calculated for each frame using CHARMM. Because the force is the negative gradient of the potential energy, and the potential energy of the restraint (*E*_res_) is calculated with the equation 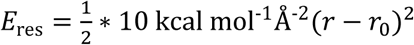, where *r* is the current location and *r*_0_ is the position of the restraint, each force component is calculated with the equation *f*_x_ = −10 *k*cal mol^-1^ Å^-2^(*x* - *x*_0_), and then converted to pN^33^. Thus, a 10 pN force applied along the x-axis would cause a displacement of 0.014 Å, since 1 kcal mol^-1^ Å^-2^ is equivalent to 69.5 pN. The magnitude of the force was then calculated as 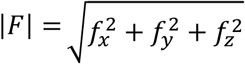. The average magnitude of the force and the backbone RMSDs at 310 K were then calculated for each stability and linker length using the WHAM equations^49^. In order to isolate the effect of co-translational folding, the quantity Δ〈|*Force*|〉 was computed as

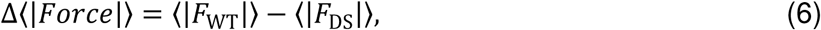

where 〈|*F*_W*T*_|〉 is the average magnitude of the force, averaged over all the saved configurations from the simulations, and 〈|*F*_DS_|〉 is the average magnitude of the force of the destabilized version of the protein at that same linker length and temperature. In this way, the force arising solely from the folding is measured.

### Temperature quench simulations

Temperature-quench simulations were performed by simulating the system at 800 K for 120 ns, then instantaneously quenching to 310 K for 1,500 ns or until the domain was folded. The time required to fold was defined by the first occurrence where the RMSD dropped below the threshold and remained below that threshold for 10^5^ timesteps or 1.5 ns. 100 trajectories were simulated for each system at each nascent-chain length. Error bars represent 95% confidence intervals computed from Bootstrapping 10,000 times.

## Supporting information

## Acknowledgements

We thank Patrick Dudas for helping create Figures 4, 5, and 7, Liam Jackson and O’Brien Lab members for a critical reading of the manuscript. We acknowledge funding from NSF grant MCB-1553291.

