## Supplementary material for "Domain topology, stability, and translation speed determine mechanical force generation on the ribosome"

### SI Methods

#### Creation of folding-incompetent domains

To create destabilized (i.e. non-folding) proteins the same procedure was used to determine an η value that resulted in a native state that was 1.81 kcal/mol higher in free energy than the unfolded state at 310 K. The domain is unfolded 95% of the time at this stability. To ensure that these destabilized proteins were not adopting partially-folded intermediates, all RMSDs were above the threshold value.

#### Rewiring of domains

Proteins 1NY8 and 1P9K were rewired to change their topology. To create these rewired mutants, all secondary structural elements in the all-atom structure were fixed in space in their crystallographic configuration, and the unstructured loops were cut and reconnected to change the order of the secondary structural elements along the primary structure. The loop residues were then minimized using the CHARMM22 force-field, during which the secondary structural elements were constrained. The resulting structure was then coarse-grained as previously described, and tuned to the same native-state stability as the wild-type domain.

#### Structural analyses

Backbone Cα RMSDs from the crystal structure were calculated in CHARMM using only those residues involved in secondary structural elements of at least 4 residues in length as defined by STRIDE[1]. For folding pathway analysis, the fraction of native contacts and the backbone RMSD were calculated for each secondary structure element. The folding thresholds for the proteins 1F0Z, 1NY8, 2IST, and 2JSO were the average RMSD values at the melting temperature (the maximum in heat capacity), while 1P9K, and its corresponding mutants, used a threshold of 8 Å due to a single strand that melts at a much lower temperature than the rest of the domain.

#### Derivation of Eq. 1

For a two-state system the free energy of stability of the folded state ($N$) relative to the unfolded state ($U$) is

$$\Delta G_{\mathrm{UN},L} = G_{\mathrm{N},L} - G_{\mathrm{U},L}, \tag{S1}$$

where $G_{\mathrm{X},L}$ is the free energy of state X at linker length $L$. We want to model the behavior of the change in force arising from folding (as defined by Eq. 6) at a fixed linker length as a function of $\Delta G_{\mathrm{UN},L}$. To do this we need to model the force arising from the wild-type protein, $\langle|F_{\mathrm{WT}}|\rangle$, and the destabilized version of it $\langle|F_{\mathrm{DS}}|\rangle$.

The force of the wild-type is determined by the forces associated with the folded and unfolded ensembles of the domain, and probabilities of populating those states. For a two state system then

$$\langle|F_{\mathrm{WT},L}|\rangle = F_{\mathrm{U},L}P(\mathrm{U}|L) + F_{\mathrm{N},L}P(\mathrm{N}|L) \tag{S2}$$

And in terms of the free energy difference between the two states

$$\langle|F_{\mathrm{WT},L}|\rangle = \frac{F_{\mathrm{U},L}e^{-G_{\mathrm{U},L}\beta} + F_{\mathrm{N},L}e^{-G_{\mathrm{N},L}\beta}}{e^{-G_{\mathrm{U},L}\beta} + e^{-G_{\mathrm{N},L}\beta}}. \tag{S3}$$

Multiplying the right-hand-side of Eq. S3 by the ratio $\frac{e^{G_{\mathrm{N},L}\beta}}{e^{G_{\mathrm{N},L}\beta}}$ yields

$$\langle|F_{\mathrm{WT},L}|\rangle = \frac{F_{\mathrm{U}} e^{\beta\Delta G_{\mathrm{UN},L}} + F_{\mathrm{N},L}}{e^{\beta\Delta G_{\mathrm{UN},L}} + 1}. \tag{S4}$$

The destabilized mutant only populates the unfolded state, so

$$\langle|F_{\mathrm{DS},L}|\rangle = F_{\mathrm{U},L}. \tag{S5}$$

Substituting Eq. S4 and S5 into Eq. 6 yields

$$\Delta\langle|Force|\rangle = \frac{F_{\mathrm{U},L} e^{\beta\Delta G_{\mathrm{UN},L}} + F_{\mathrm{N},L}}{e^{\beta\Delta G_{\mathrm{UN},L}} + 1} - F_{\mathrm{U},L}. \tag{S6}$$

Through the following series of algebraic rearrangements

$$\Delta\langle|Force|\rangle\left[e^{\beta\Delta G_{\mathrm{UN},L}} + 1\right] = F_{\mathrm{U},L} e^{\beta\Delta G_{\mathrm{UN},L}} + F_{\mathrm{N},L} - F_{\mathrm{U},L}\left[e^{\beta\Delta G_{\mathrm{UN},L}} + 1\right] \tag{S7}$$

$$\Delta\langle|Force|\rangle\left[e^{\beta\Delta G_{\mathrm{UN},L}} + 1\right] = F_{\mathrm{U},L} e^{\beta\Delta G_{\mathrm{UN},L}} + F_{\mathrm{N},L} - F_{\mathrm{U},L} e^{\beta\Delta G_{\mathrm{UN},L}} - F_{\mathrm{U},L} \tag{S8}$$

$$\Delta\langle|Force|\rangle\left[e^{\beta\Delta G_{\mathrm{UN},L}} + 1\right] = \left[F_{\mathrm{N},L} - F_{\mathrm{U},L}\right], \tag{S9}$$

we get the result that

$$\Delta\langle|Force|\rangle = \frac{\left[F_{\mathrm{N},L} - F_{\mathrm{U},L}\right]}{e^{\beta\Delta G_{\mathrm{UN},L}} + 1} \tag{1}$$

for a two state system. For a system with $I$ intermediate states, this equation may be expanded to

$$\Delta\langle|Force|\rangle = \frac{[F_{\mathrm{N}} - F_{\mathrm{U}}] + \sum_{i\in I} e^{\beta\Delta G_{i\mathrm{N},L}}(F_i - F_{\mathrm{U}})}{1 + e^{\beta\Delta G_{\mathrm{UN},L}} + \sum_{i\in I} e^{\beta\Delta G_{i\mathrm{N},L}}}. \tag{S10}$$

**Derivation of Eq. 2**

Equation 1 allows us to predict the force as a function of free energy at a given linker length. As the protein becomes more likely to fold at longer linker lengths, the increasingly negative free energy prevents the force from returning to zero at extended linker lengths (Fig. S3), so we write the free energy as a function of $L$ ($\Delta G_{\mathrm{UN}}(L)$). Force generation and transmission back to the PTC requires three factors—the domain must be folded, in contact with the ribosome, and there must be minimal slack in the linker. Thus, to account for these, we multiply Eq. 1 by the probability that the end-to-end distance of the linker ($L_{\mathrm{ee}}$) divided by the contour length of the linker ($L_{\mathrm{C}}$) is greater than a threshold value ($x > 0.73$) when the domain is folded ($N$) and touching the ribosome ($T$) at a given linker length ($L$), which gives

$$\Delta\langle|Force|\rangle = \frac{[F_{\mathrm{N}} - F_{\mathrm{U}}]\, P\left(\frac{L_{\mathrm{ee}}}{L_{\mathrm{C}}} > x | L, \mathrm{N}, T\right)}{e^{\beta\Delta G_{\mathrm{UN}}(L)} + 1}. \tag{S11}$$

To identify a threshold value to determine if there is slack in the linker, we used the generalized bead rod polymer model[2] to estimate the force required to stretch a generic polymer in an inert cylinder, and then identified low-, intermediate-, and high-force regimes in the plot of force versus normalized end-to-end distance, with our threshold value (0.73) coming from the boundary of the intermediate and high-force regimes (Fig. 9). However, Eq. S11 only applies when the system is in equilibrium, and to explain the effects of translation speed, it must be extended to cover both equilibrium and non-equilibrium systems. As the quantity $\frac{1}{e^{\beta\Delta G_{\mathrm{UN}}(L)} + 1}$ is equivalent to $P(\mathrm{N}|L)$, and $P(\mathrm{N}|L) = \frac{\omega_{\mathrm{A}}(L+1)P(\mathrm{N}|L-1) + k_{\mathrm{N}}(L)}{k_{\mathrm{N}}(L) + k_{\mathrm{U}}(L) + \omega_{\mathrm{A}}(L+1)}$ [3], where $k_{\mathrm{N}}(L)$ and $k_{\mathrm{U}}(L)$ are the folding and unfolding rates and $\omega_A(L+1)$ is the rate at which the next codon is translated, we can write Eq. S11 as

$$\Delta\langle|Force|\rangle = [F_{\mathrm{N}} - F_{\mathrm{U}}] P\left(\frac{L_{\mathrm{ee}}}{L_{\mathrm{C}}} > 0.73 \middle| L, \mathrm{N}, T\right) \frac{\omega_{\mathrm{A}}(L+1)P(\mathrm{N}|L-1) + k_{\mathrm{N}}(L)}{k_{\mathrm{N}}(L) + k_{\mathrm{U}}(L) + \omega_{\mathrm{A}}(L+1)}. \tag{2}$$

As above, equation 2 can be expanded to include $I$ intermediate states.

For this statistical mechanical model, the domain was defined to be touching the ribosome surface if at least one residue of the domain was within 8.8 Å of any ribosomal interaction site, which was selected because it encompasses 90% of native contacts in the five domains used. For a measure of slack in the linker, the distance between the first and last residues was measured and normalized by the contour length of the linker, which equals 3.8 Å(N-1), where N is the number of Cα sites in the linker.

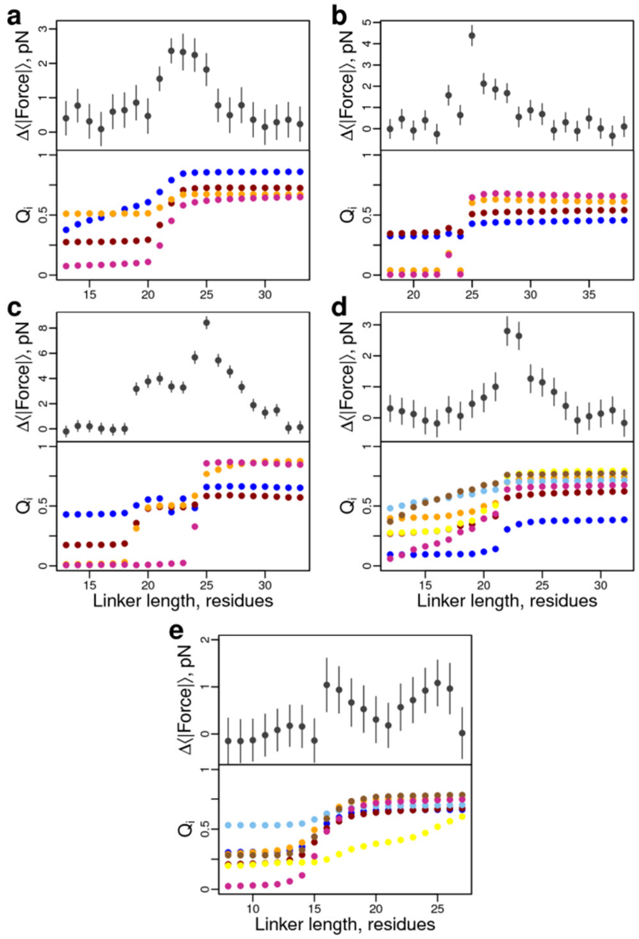


**Figure S1. Fraction of native contacts per secondary structural element reveals cooperative and non-cooperative folding transitions give rise to pulling forces.** Gray circles correspond to the average pulling force at the C-terminal nascent chain residue as a function of linker length for the five different proteins: (**a**) 1F0Z, (**b**) 2IST, (**c**) 1P9K, (**d**) 2JSO, and (**e**) 1NY8. Colored circles (lower graphs) correspond to the fraction of native contacts formed for each secondary structural element (α-helix or β-strand). 1P9K and 1NY8 exhibit non-cooperative transitions, meaning a partially folded intermediate is first populated before the native state is fully formed at longer linker lengths. Comparing the force curves to the native contact curves reveals folding (or partially folding) transitions correlate with large pulling forces. Error bars represent 95% confidence intervals about the mean calculated from Block Averaging.

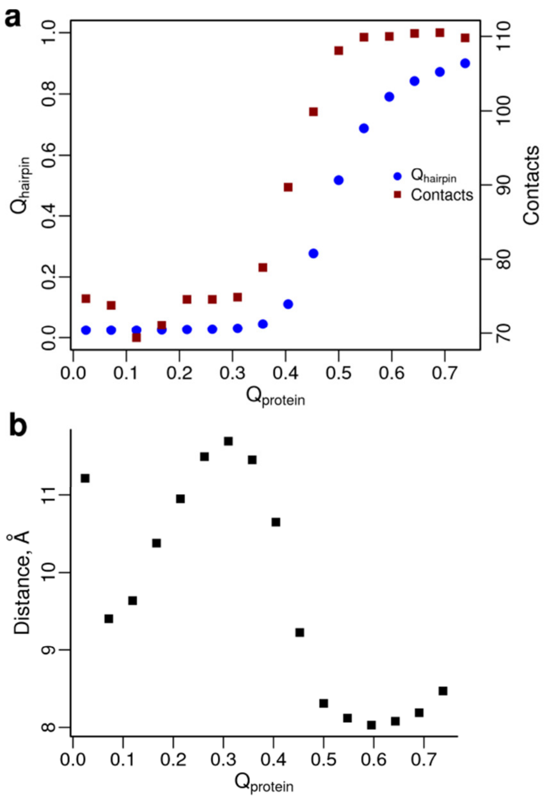


**Figure S2. Folding of a C-terminal β-hairpin causes a domain to press against the ribosome surface. (a)** Blue circles indicate the fraction of native intra-hairpin contacts formed as a function of the total native contacts in the domain, and red squares indicate the number of contacts between the domain from protein 1P9K and the ribosome surface at 310 K and a linker length of 25 residues. The correlation between the two data sets reveals that hairpin formation and increase in contacts between the domain and ribosome are concomitant. **(b)** Same system as **(a)**, except plotted is the average minimum distance between residue 66 (the most N-terminal residue in the β-hairpin) and the ribosome surface as a function of the fraction of native contacts in the full domain. The trend indicates that as the hairpin folds the more N-terminal strand flips back towards the ribosome surface.Note that the average $Q_{protein}$ is 0.65 at 310 K. Therefore, we set the upper limit of the range on the X-axis to 0.1 above this value in panels **(a)** and **(b)**.

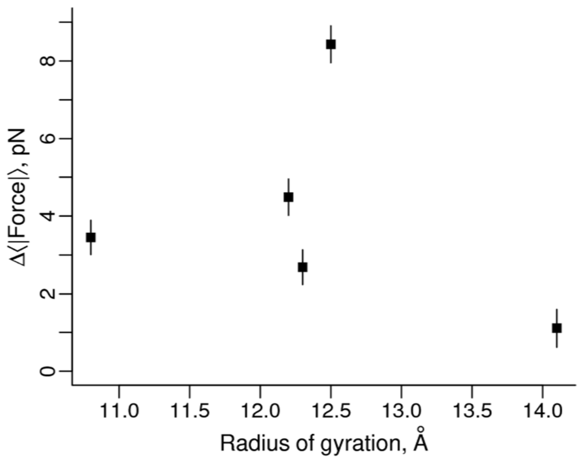


**Figure S3. Domain size has no apparent effect on the magnitude of the pulling force.** The pulling force versus the radius of gyration as a measure of domain size for the five domains. No statistically significant trend is observed. Error bars represent 95% confidence intervals about the mean calculated from Block Averaging.

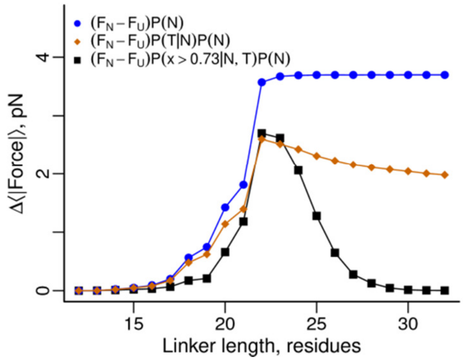


**Figure S4. Folding, contact with the ribosome, and no slack in the linker are all essential factors for accurately describing force profiles.** Black squares represent the pulling force model (Eq. 2) for protein 2JSO, which incorporates the probability the linker is stretched at least 73% of its contour length given that the domain is folded and in contact with the ribosome. The x in the legend represents the ratio $\frac{\boldsymbol{L}_{\mathrm{ee}}}{\boldsymbol{L}_{\mathrm{C}}}$, and $\boldsymbol{P}(\mathrm{N})$ is equal to $\frac{\mathbf{1}}{\mathbf{1}+\boldsymbol{e}^{\boldsymbol{\beta}\Delta\boldsymbol{G}_{\mathrm{UN}}(\boldsymbol{L})}}$. Here we systematically examine how each model feature contributes to the shape of the force profile while keeping $[\boldsymbol{F}_{\mathrm{N}} - \boldsymbol{F}_{\mathrm{U}}]$ fixed. Blue circles correspond to only accounting for the probability the domain is folded. Orange diamonds are the force when the population of folded domains that are touching the ribosome surface is accounted for (through the product of $\boldsymbol{P}(\boldsymbol{T}|\mathrm{N})\boldsymbol{P}(\mathrm{N})$). It is only when additionally accounting for slack in the linker that the right force profile shape emerges (black squares).

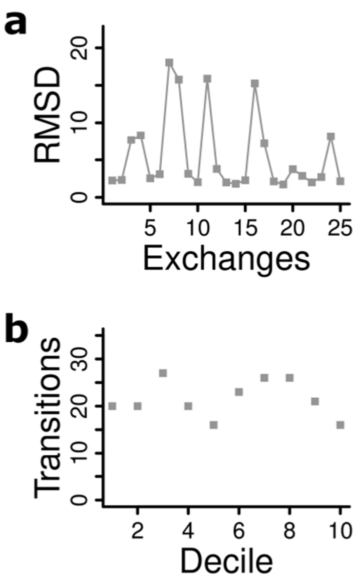


**Figure S5**.**Necessary criteria for replica exchange simulations. (a)** RMSD versus number of replica exchanges illustrates the extensive sampling of folded and unfolded states achieved using replica-exchange simulations. Data from 1F0Z on the ribosome at 326 K and a linker of 25 residues. **(b)** Number of exchanges between high- and low-temperature windows per decile of a 300,000 replica-exchange simulation of1F0Z on the ribosome. The lack of a systematic trend in the data is a necessary but not sufficient condition for equilibrium sampling.

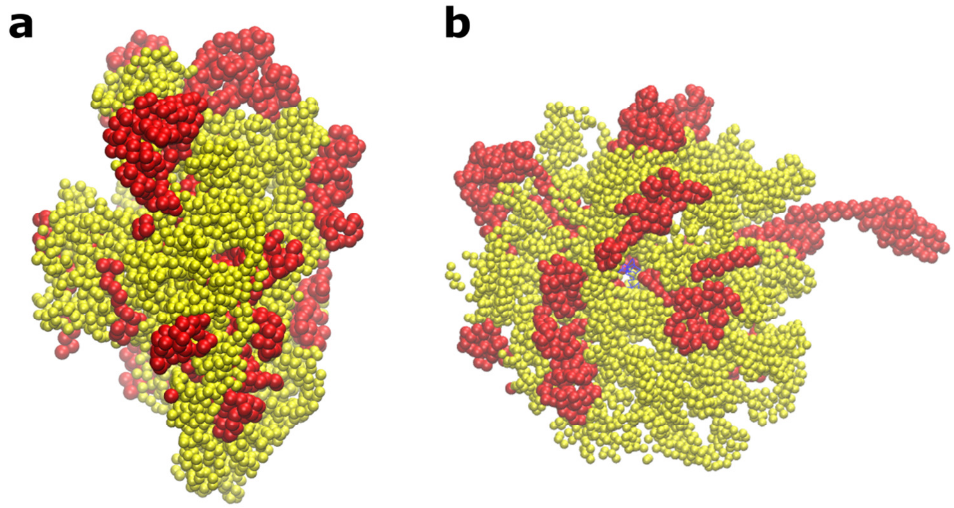


**Figure S6. A ribosome cutout was used to reduce simulation costs.** (**a**) Side and (**b**) top views of the ribosome cutout, with yellow representing RNA, red representing ribosomal proteins, and the nascent chain in the exit tunnel in blue (b). The ribosome was trimmed to retain all residues within 10 Å of the exit tunnel and all surface residues accessible to the largest unfolded domain at a linker length of 50 residues.

| Protein | PDB ID | Structural class | Residues | $\eta^*$ | Closest $\eta$ | $Q_{eq}$ | $Q_{310}$ | $Q_{kin} = \frac{Q_{eq}+Q_{310}}{2}$ | $\langle\tau_N^{sim}\rangle$ (ns) | $\langle\tau_N^{exp}\rangle$ (s) | $\alpha$, $\langle\tau_N^{exp}\rangle/\langle\tau_N^{sim}\rangle$ |
|---|---|---|---|---|---|---|---|---|---|---|---|
| EC298 | 1RYK | α | 69 | 1.080 | 1.100 | 0.67 | 0.98 | 0.83 | 8.38 | 0.00011[4] | 13,591 |
| λ-repressor | 1LMB | α | 80 | 1.163 | 1.200 | 0.53 | 0.83 | 0.68 | 26.54 | 0.00003[4] | 1,170 |
| bACBP | 2ABD | α | 86 | 1.304 | 1.350 | 0.62 | 0.94 | 0.78 | 11.82 | 0.00095[4] | 80,574 |
| IM7 | 1CEI | α | 85 | 0.998 | 1.000 | 0.62 | 0.76 | 0.69 | 49.52 | 0.00075[4] | 15,070 |
| IM9 | 1IMQ | α | 86 | 1.298 | 1.300 | 0.51 | 0.78 | 0.64 | 6.44 | 0.00066[4] | 102,158 |
| Cytochrome-256 | 256B | α | 106 | 1.174 | 1.150 | 0.57 | 0.87 | 0.72 | 18.20 | 0.00125[5] | 68,681 |
| ABP1 SH3 | 1JO8 | β | 58 | 1.628 | 1.600 | 0.77 | 1.0 | 0.89 | 51.62 | 0.08547[4] | 1,655,755 |
| Fyn SH3 | 1SHF | β | 59 | 1.467 | 1.500 | 0.51 | 0.98 | 0.75 | 7.15 | 0.01060[4] | 1,483,140 |
| CspB | 1C9O | β | 66 | 1.298 | 1.300 | 0.48 | 0.74 | 0.61 | 18.02 | 0.00073[4] | 40,506 |
| CspA | 1MJC | β | 69 | 1.445 | 1.450 | 0.51 | 0.78 | 0.64 | 18.08 | 0.00503[6] | 278,208 |
| Tenascin | 1TEN | β | 89 | 1.457 | 1.500 | 0.39 | 0.82 | 0.61 | 12.04 | 0.16611[4] | 13,796,757 |
| Twitchin | 1WIU | β | 93 | 1.357 | 1.400 | 0.47 | 0.82 | 0.65 | 18.63 | 0.66667[4] | 35,784,577 |
| Hpr | 1POH | α/β | 85 | 1.164 | 1.150 | 0.48 | 0.86 | 0.67 | 68.98 | 0.06711[7] | 972,891 |
| Urm1 | 2QJL | α/β | 99 | 1.075 | 1.100 | 0.47 | 0.86 | 0.66 | 284.55 | 0.07576[4] | 266,236 |
| Src SH2 | 1SPR | α/β | 102 | 1.298 | 1.300 | 0.67 | 0.94 | 0.81 | 6.57 | 0.00016[4] | 24,353 |
| Azurin (apo) | 1E65 | α/β | 128 | 1.163 | 1.200 | 0.70 | 0.90 | 0.80 | 203.40 | 0.00735[4] | 36,136 |
| CheY | 3CHY | α/β | 128 | 1.001 | 1.000 | 0.46 | 0.75 | 0.61 | 96.55 | 0.02653[8] | 274,780 |
| Ribonuclease H | 2RN2 | α/β | 155 | 1.047 | 1.100 | 0.22 | 0.59 | 0.41 | 81.80 | 1.35135[9] | 16,520,171 |
| | | | | | | | | | | | $\langle\alpha\rangle = 3{,}967{,}486$ |

**Table S1: Kinetic parameters for the 18 proteins in the training set.** Results of temperature quenching for all 18 training set proteins. Note that a kinetic definition of the $Q$ threshold, $Q_{\text{kin}} = \frac{Q_{\text{eq}}+Q_{310}}{2}$, is used in this analysis. The acceleration factor, α, is the fold acceleration of protein folding in our simulations compared to experiment.

| |
|---|
| N-terminus-AVQLALAALISALEKEVVILLALVKALGALLLLLAALAALAAIDALELVA-C terminus |

**Table S2. Unstructured linker sequence used in all simulations[10].**

| Protein | PDB ID | Experimental Parameters | | REX input and output parameters | | | | Linear regression parameters | | | |
|---|---|---|---|---|---|---|---|---|---|---|---|
| | | $T_{exp}$ (K) | $\Delta G_{UN}^{exp}$ (kcal/mol) | η | $T_M$ (K) | $Q_{eq}$ | $\Delta G_{UN}^{sim}(T_{exp})$ (kcal/mol) | $m$ | $b$ | $R^2$ | $\eta^*$ |
| EC298 | 1RJK | 298 | -2.72[4] | 1.000<br>1.050<br>1.100<br>1.200 | 302.6<br>311.2<br>319.0<br>335.0 | 0.67<br>0.67<br>0.67<br>0.65 | -0.798<br>-2.10<br>-3.17<br>-5.53 | -23.456 | 22.609 | 0.99927 | 1.080 |
| λ-repressor | 1LMB | 298 | -4.25[4] | 1.200<br>1.300<br>1.400 | 321.0<br>336.4<br>351.6 | 0.53<br>0.55<br>0.51 | -5.13<br>-6.77<br>-9.17 | -20.230 | 19.278 | 0.98838 | 1.163 |
| bACBP | 2ABD | 298 | -6.40[4] | 1.200<br>1.255<br>1.350 | 316.4<br>323.8<br>336.8 | 0.66<br>0.68<br>0.62 | -4.26<br>-5.09<br>-7.49 | -21.941 | 22.216 | 0.98517 | 1.304 |
| IM7 | 1CEI | 298 | -2.88[4] | 1.000<br>1.150<br>1.200 | 313.6<br>341.2<br>351.6 | 0.62<br>0.59<br>0.58 | -2.96<br>-5.22<br>-6.42 | -16.745 | 13.832 | 0.98901 | 0.998 |
| IM9 | 1IMQ | 298 | -6.81[4] | 1.050<br>1.100<br>1.150<br>1.300<br>1.350<br>1.400 | 307.0<br>315.0<br>322.8<br>345.8<br>353.8<br>362.2 | 0.53<br>0.52<br>0.52<br>0.51<br>0.50<br>0.48 | -1.90<br>-3.39<br>-4.42<br>-6.69<br>-7.36<br>-9.05 | -18.480 | 17.167 | 0.98347 | 1.298 |
| Cytochrome-256b | 256B | 298 | -6.60[11] | 1.000<br>1.150<br>1.300<br>1.600 | 301.8<br>321.8<br>341.0<br>378.4 | 0.60<br>0.57<br>0.56<br>0.55 | -1.18<br>-6.46<br>-10.3<br>-18.3 | -28.079 | 26.378 | 0.99558 | 1.174 |

**Table S3: Replica exchange results for α-helical proteins.** Replica exchange results for all α-helical proteins are shown. The values of $T_{\mathrm{M}}$, $\mathrm{Q}_{\mathrm{eq}}$, and $\Delta G_{\mathrm{UN}}^{\mathrm{sim}}(T_{\mathrm{exp}})$ were calculated using the WHAM equation. All free energies, *m*, and *b* are in units of kcal/mol.

| Protein | PDB ID | Experimental Parameters $\mathbf{T_{exp}}$ (K) | $\Delta G_{UN}^{exp}$ (kcal/mol) | REX input and output parameters η | $\mathbf{T_M}$ (K) | $\mathbf{Q_{eq}}$ | $\Delta G_{UN}^{sim}(T_{exp})$ (kcal/mol) | Linear regression parameters $m$ | $b$ | $R^2$ | η* |
|---|---|---|---|---|---|---|---|---|---|---|---|
| ABP1 SH3 | 1JO8 | 298 | -3.07[4] | 1.350<br>1.400<br>1.500<br>1.600 | 316.6<br>324.0<br>338.0<br>352.6 | 0.77<br>0.77<br>0.77<br>0.77 | -1.75<br>-2.06<br>-2.52<br>-2.91 | -4.590 | 4.403 | 0.99358 | 1.628 |
| Fyn SH3 | 1SHF | 293 | -6.68[4] | 1.350<br>1.400<br>1.500 | 333.2<br>339.8<br>355.4 | 0.51<br>0.54<br>0.51 | -5.48<br>-5.76<br>-7.10 | -11.149 | 9.680 | 0.97134 | 1.467 |
| CspB Bc | 1C9O | 298 | -1.81[4] | 1.300<br>1.400<br>1.500<br>1.600 | 311.4<br>326.2<br>340.6<br>354.4 | 0.48<br>0.47<br>0.47<br>0.47 | -1.69<br>-3.38<br>-4.71<br>-5.83 | -13.749 | 16.035 | 0.99129 | 1.298 |
| CspA | 1MJC | 298 | -3.00[11] | 1.450<br>1.500<br>1.600 | 317.4<br>324.0<br>337.2 | 0.51<br>0.50<br>0.51 | -3.02<br>-3.93<br>-5.33 | -15.226 | 18.998 | 0.99500 | 1.445 |
| Tenascin | 1TEN | 298 | -6.71[4] | 1.400<br>1.500<br>1.600 | 329.0<br>343.8<br>360.4 | 0.40<br>0.39<br>0.39 | -5.16<br>-7.81<br>-10.8 | -28.220 | 34.408 | 0.99883 | 1.457 |
| Twitchin | 1WIU | 293 | -5.00[4] | 1.200<br>1.300<br>1.400 | 301.4<br>314.8<br>329.0 | 0.49<br>0.49<br>0.47 | -1.73<br>-4.06<br>-5.77 | -20.195 | 22.402 | 0.99238 | 1.357 |

**Table S4: Replica exchange results for β-sheet proteins.** Replica exchange results for all β-sheet proteins are shown. The values of $T_M$, $Q_{eq}$, and $\Delta G_{UN}^{sim}(T_{exp})$ were calculated using the WHAM equation. All free energies, *m*, and *b* are in units of kcal/mol.

| Protein | PDB ID | Experimental Parameters | | REX output parameters | | | | Linear regression parameters | | | |
|---|---|---|---|---|---|---|---|---|---|---|---|
| | | $T_{exp}$ (K) | $\Delta G_{UN}^{exp}$ (kcal/mol) | $\eta$ | $T_M$ (K) | $Q_{eq}$ | $\Delta G_{UN}^{sim}(T_{exp})$ (kcal/mol) | $m$ | $b$ | $R^2$ | $\eta^*$ |
| Hpr | 1POH | 298 | -5.46[4] | 1.050 | 308.6 | 0.51 | -2.79 | -21.377 | 19.419 | 0.94639 | 1.164 |
| | | | | 1.150 | 323.0 | 0.48 | -5.77 | | | | |
| | | | | 1.200 | 330.2 | 0.49 | -6.03 | | | | |
| | | | | 1.230 | 335.6 | 0.49 | -6.71 | | | | |
| Urm1 | 2QJL | 298 | -3.48[4] | 1.100 | 312.6 | 0.47 | -3.74 | -22.836 | 21.078 | 0.98269 | 1.075 |
| | | | | 1.200 | 331.0 | 0.48 | -6.56 | | | | |
| | | | | 1.300 | 348.8 | 0.47 | -9.04 | | | | |
| | | | | 1.400 | 366.2 | 0.49 | -10.5 | | | | |
| Src SH2 | 1SPR | 298 | -7.24[4] | 1.100 | 317.6 | 0.72 | -3.72 | -20.051 | 18.778 | 0.91770 | 1.298 |
| | | | | 1.190 | 324.8 | 0.71 | -4.53 | | | | |
| | | | | 1.200 | 329.0 | 0.73 | -5.01 | | | | |
| | | | | 1.300 | 347.6 | 0.67 | -7.67 | | | | |
| Azurin (apo) | 1E65 | 298 | -5.29[4] | 1.050 | 303.4 | 0.69 | -2.55 | -25.190 | 23.997 | 0.98516 | 1.163 |
| | | | | 1.100 | 312.2 | 0.69 | -4.10 | | | | |
| | | | | 1.200 | 328.0 | 0.70 | -5.49 | | | | |
| | | | | 1.300 | 345.8 | 0.64 | -8.72 | | | | |
| | | | | 1.400 | 362.2 | 0.61 | -11.5 | | | | |
| CheY | 3CHY | 298 | -5.60[11] | 1.000 | 315.2 | 0.46 | -5.48 | -58.105 | 52.554 | 0.99949 | 1.001 |
| | | | | 1.100 | 335.6 | 0.46 | -11.5 | | | | |
| | | | | 1.200 | 354.2 | 0.44 | -17.1 | | | | |
| Ribonuclease H | 2RN2 | 298 | -7.20[11] | 1.100 | 332.4 | 0.22 | -9.19 | -38.735 | 33.369 | 0.99945 | 1.047 |
| | | | | 1.200 | 348.4 | 0.22 | -13.2 | | | | |
| | | | | 1.300 | 366.0 | 0.22 | -16.9 | | | | |
| Dihydrofolate reductase | 1RX4 | 298 | -6.37[11] | 0.930 | 305.2 | 0.51 | -3.01 | -29.666 | 24.797 | 0.98363 | 1.051 |
| | | | | 1.000 | 315.0 | 0.55 | -4.49 | | | | |
| | | | | 1.100 | 331.4 | 0.55 | -7.99 | | | | |

**Table S5: Replica exchange results for α/β proteins.** Replica exchange results for all α/β proteins are shown. The values of $T_M$, $Q_{eq}$, and $\Delta G_{UN}^{sim}(T_{exp})$ were calculated using the WHAM equation. All free energies, *m*, and *b* are in units of kcal/mol.
